# The contribution of short tandem repeats to splicing variation in the human prefrontal cortex

**DOI:** 10.64898/2026.05.04.721407

**Authors:** Yang Li, Jonathan Margoliash, Alon Goren, Melissa Gymrek

**Affiliations:** Department of Medicine, University of California San Diego, La Jolla, CA, USA; Department of Computer Science and Engineering, University of California San Diego, La Jolla, CA, USA; Department of Pediatrics, University of California San Diego, La Jolla, CA, USA

**Keywords:** Short tandem repeats (STRs), Alternative splicing, Splicing quantitative trait loci (sQTLs), Schizophrenia, RNA-binding proteins (RBPs), Colocalization analysis, Fine-mapping analysis, Human brain

## Abstract

**Background:** Splicing disruption has been implicated in a range of heritable phenotypes, including numerous psychiatric and neurological disorders. Recent studies have identified thousands of common genetic variants impacting splicing in brain and other tissues, but have focused largely on single nucleotide polymorphisms or short indels. Despite growing evidence that genetic variation at short tandem repeats (STRs) influences splicing, large-scale studies of STR-mediated splicing in brain have been limited by low sample sizes of available RNA-seq data or exclusion of certain classes of STRs, such as homopolymers which account for around half of all STRs.

**Results:** In this study, we leveraged deep RNA-seq and SNP array data from 336 human dorsolateral prefrontal cortex (DLPFC) samples collected by the Human Brain Collection Core (HBCC). We imputed 445,720 STRs into available genotype data and identified 51,343 unique STRs for which copy number is significantly associated with one or multiple alternative splicing events of nearby genes (spliceSTRs). We prioritized and characterized candidate causal spliceSTRs using three orthogonal fine-mapping strategies which identified 1,313 high-confidence fine-mapped spliceSTRs. Our analyses revealed strong associations between copy number of certain repeat units and binding of specific RNA-binding proteins (RBPs), including a previously known relationship between HNRNPL and *AC* repeat length, suggesting that the functional impact of some spliceSTRs may be mediated through their binding affinity for RBPs. Finally, co-localization analyses using summary statistics from genome-wide association studies (GWAS) for 6 brain-related disorders identified multiple signals that may be driven by spliceSTRs, including a previously identified *GT*_n_ repeat that is a spliceSTR for *PLEKHA1* associated with Alzheimer’s disease as well as a newly identified *AGG*_n_ spliceSTR in *SEPTIN3* co-localized with schizophrenia.

**Conclusions:** Together, our findings highlight the role of STRs in regulating alternative splicing in the human brain, suggest a general relationship between STR polymorphism and RBP-mediated splicing events, and identify specific spliceSTRs detected by co-localization analysis as candidate causal variants for brain-related disorders.

## Background

Alternative splicing is a regulatory mechanism by which a single gene can produce different isoforms, greatly expanding the diversity of the transcriptome and proteome. This process is tightly regulated and occurs in approximately 90–95% of human genes, playing a critical role in development, cell differentiation^1,2^, and tissue-specific gene expression^3^. Regulation of alternative splicing is highly dependent on the underlying nucleotide sequence in *cis* and is recognized as a primary link between genetic variation and diseases^4^ ranging from neurological disorders to cancer^5,6^. Dysregulation of alternative splicing is particularly known to be involved in many heritable brain-related disorders including spinal muscular atrophy (SMA)^7–9^, myotonic dystrophy (DM1 and DM2)^10–12^, and schizophrenia^13–15^.

Recent splicing quantitative trait locus (sQTL) analyses have identified tens of thousands of variants associated with alternative splicing variation across diverse human tissues^16,17^ including brain^18^. Yet, most of those studies have focused on single-nucleotide polymorphisms (SNPs) and short insertions or deletions (indels). Short tandem repeats (STRs), consisting of repeated units of 1-6bp, comprise an additional large source of genetic variation^19,20^. Despite increasing evidence of a role for STR variation in a range of human traits, including gene regulation^21–26^ and other complex traits^27^, they are often excluded from genome-wide studies since they are challenging to genotype and require modified association testing paradigms.

Emerging evidence suggests that variation in copy number of STRs can modulate alternative splicing, contributing to both normal variation in alternative splicing as well as to the pathogenesis of a wide range of diseases. For example, length of a (*TG*)_n_ repeat is associated with exon 9 inclusion of *CFTR*. This splicing event can influence the penetrance of other pathogenic mutations in the *CFTR* gene which are implicated in nonclassic cystic fibrosis and male infertility^28,29^. In other cases, large repeat expansions have been shown to cause global changes in alternative splicing by sequestering splicing factors. For example, a (*GGGGCC*)_n_ expansion that leads to aberrant splicing of *C9orf72*, has been identified as a major cause of both dominant frontotemporal dementia (FTD) and amyotrophic lateral sclerosis (ALS)^30,31^. This (*GGGGCC*)_n_ expansion has been shown to sequester the RNA binding proteins (RBPs) TDP-43 and HNRNPH, leading to their mis-localization and downstream splicing defects^32–36^. In another example, expansion of (*CTG*)_n_/(*CCTG*)_n_ repeats implicated in myotonic dystrophy have been shown to sequester the splicing factor MBNL1^37,38^.

Two recent studies have investigated the genome-wide role of STR variation in alternative splicing. Hamanaka *et al.* analyzed 49 tissues from the Genotype-Tissue Expression (GTEx) dataset^17^, which revealed that repeat-associated splicing variation in the brain was distinct from that in non-brain tissues^39^. Yet, the small sample size, shorter read lengths, and lower sequencing depth of the GTEx brain RNA-seq datasets reduced power to detect associations in those tissues (discussed below). While the second study, Cui *et al.*^40^, identified a higher number of STR–splicing associations in brain tissues with a larger sample size, it excluded homopolymers, which account for over 50% of all STRs. Importantly, despite thousands of STR-splice event associations being identified, the mechanisms by which most of these variants act remains poorly understood.

Here, we systematically analyzed the contribution of STRs to splicing variation using RNA-seq data from 336 dorsolateral prefrontal cortex (DLPFC) samples collected by the Human Brain Collection Core (HBCC) to identify STRs for which repeat lengths are associated with nearby splicing events (spliceSTRs). We performed rigorous fine-mapping of STR and SNP sQTLs using three orthogonal approaches and characterized the resulting candidate causal spliceSTRs to identify interaction with RBPs as a potential mechanism driving a subset of spliceSTRs. Finally, we integrated the spliceSTRs we identified with summary statistics from genome-wide association studies (GWAS) to identify signals for brain-related disorders that are co-localized with fine-mapped spliceSTRs. Overall, in addition to our findings, our catalog of spliceSTRs will serve as a resource to support future investigations into complex traits and genetic variants mediating splicing variation.

## Results

### Identification of STRs associated with splicing in human dorsolateral prefrontal cortex

We obtained RNA-seq data derived from the dorsolateral prefrontal cortex for 364 samples from the HBCC^41^, comprising 209 controls and 155 cases (65 bipolar and 90 schizophrenia). Of these, we focused on 336 samples (194 controls and 142 cases) for which both SNP genotypes and high-quality RNA-seq data are available. We used RNA-seq reads to estimate exon splicing levels (percent spliced in; PSI) using rMATs^42^, which quantifies PSI for multiple classes of alternative splicing events: skipped exon (SE), mutually exclusive exon (MXE), alternative 3’ (A3SS) or 5’ (A5SS) site, and retained intron (RI) (**Methods**, **Fig. S1**). In parallel we used our published SNP-STR reference haplotype panel^43^ to impute 445,720 autosomal STRs (148,229 non-homopolymer and 297,491 homopolymers) into phased SNP genotype data for each sample (**Fig. 1a**). Overall, we found that homopolymers showed similar or greater imputation quality scores compared to non-homopolymer STRs included in our panel (**Fig. S2**).

**Figure 1.**
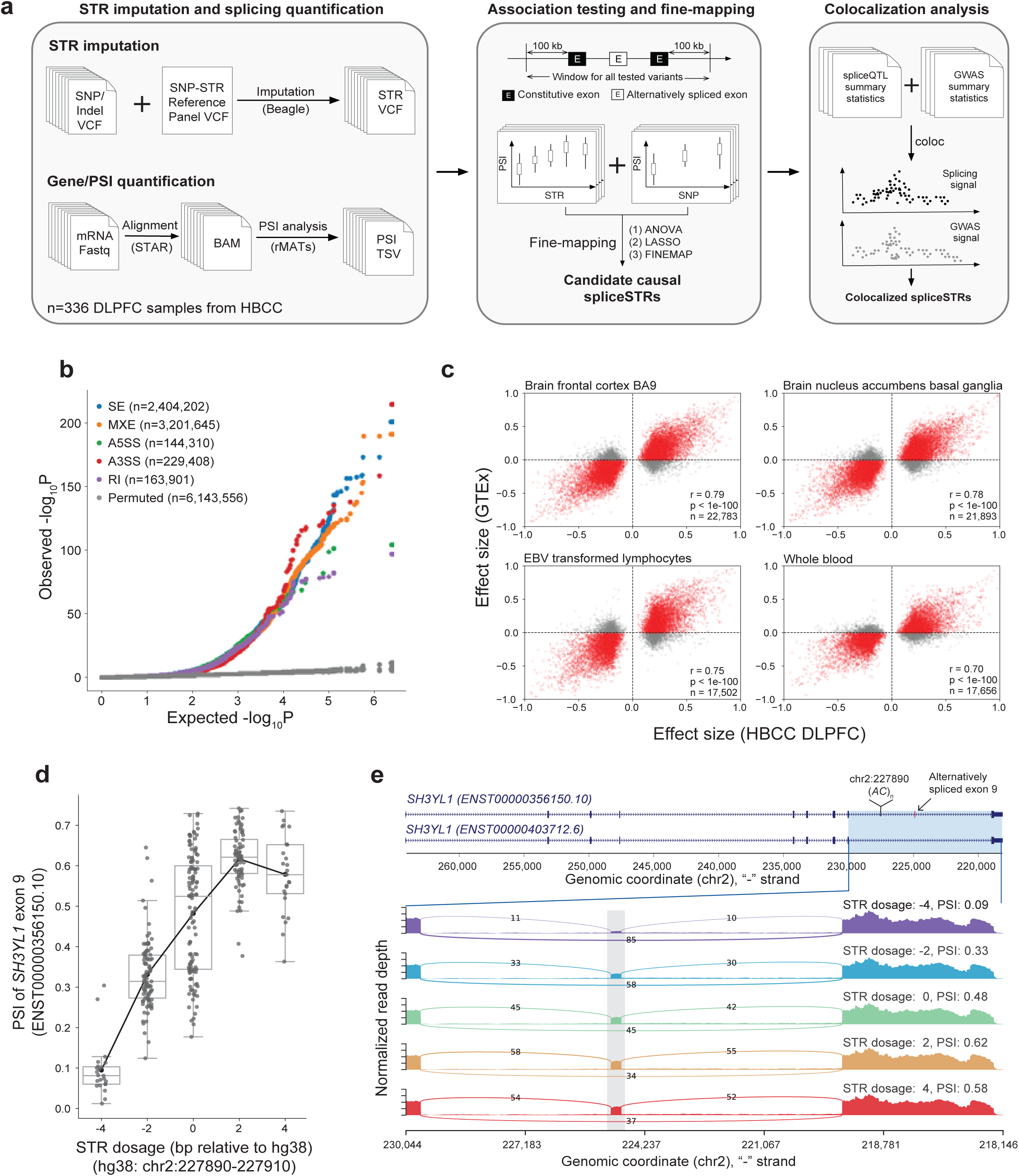
Identification of STRs associated with splicing in the human dorsolateral prefrontal cortex. **(a) Schematic overview of this study.** We used imputed STR genotypes and available RNA-seq data in HBCC to identify STRs whose lengths are associated with splicing events (spliceSTRs), followed by fine-mapping and co-localization analysis. **(b) SpliceSTR association results.** The quantile-quantile plot compares observed -log_10_ *P*-values for each STR-splice event test (*y*-axis) versus the expected uniform distribution (*x*-axis) for each type of splicing event. Gray dots denote permutation controls. **(c) Comparison of spliceSTR effect sizes across cohorts.** The *x*-axis of each plot represents effect sizes measured in HBCC. The *y*-axis of each plot represents effect sizes measured in GTEx. Only spliceSTRs reaching FDR<5% in HBCC are included. **(d) Example spliceSTR.** The *x*-axis represents the STR dosage (in bp relative to hg38) of the STR located at chr2:227890 - 227910 (hg38), which is within an intron of *SH3YL1*. The *y*-axis represents the PSI of exon 9 (ENST00000356150.10). Each dot represents an individual. Box plots summarize distributions of PSI values. Horizontal lines show median values, boxes span from the 25th percentile (Q1) to the 75th percentile (Q3). Whiskers extend to Q1 - 1.5 × IQR (bottom) and Q3 + 1.5 × IQR (top), where IQR is the interquartile range (Q3– Q1). The black line shows the mean expression for each x-axis value. **(e) Sashimi plot showing read-level evidence for the *SH3YL1* spliceSTR.** The exon structure of two representative *SH3YL1* transcripts, the location of the spliceSTR, and exons involved in the splice event (pink box) are shown in the top panel. Samples were grouped by STR dosage. The alternatively spliced exon is marked by a grey box. The number of reads supporting the skipped exon decreases with the STR dosage. Intronic regions are displayed at one-fifth the scale of exonic regions. Coordinates are in descending order since annotations are shown for the reverse strand relative to hg38. Sashimi plots were generated using rmats2sashimiplot (v2.0.4).

To detect STRs whose repeat lengths are associated with splicing, we identified all STRs within 100 kilobases (kb) of the two constitutive exons for each splicing event. For each STR-splice event pair, we tested for association between STR dosage (encoded as the sum of the best guess imputed allele length on both chromosomes) and exon splicing levels (PSI values), using a linear model with genotype principal components and PEER factors included as covariates (**Methods**). In total, we performed 6,143,556 association tests considering 255,285 unique STRs and 147,500 unique splicing events (**Supplementary Dataset 1**). We identified a total of 146,969 STR-splice event pairs (spliceSTRs) at a false discovery rate (FDR) of 5% (**Fig. 1b**, **Fig. S3**), corresponding to 51,343 unique STRs and 25,230 splicing events.

We next tested for replication of significant spliceSTRs identified in HBCC in four different tissues from the GTEx dataset^17^ (two brain tissues, whole blood, and EBV transformed lymphocytes), for which both high-coverage whole-genome sequencing (WGS) and RNA-seq data are available. Instead of relying on imputation, we directly genotyped STRs using HipSTR^19^ from the available WGS data in the GTEx dataset. Due to differences in data characteristics, including lower sample size, shorter read length, lower coverage (**Fig. S4a**), and high PCR error rates in the GTEx data (**Fig. S4b**) leading to reduced quality of STR genotypes, only a subset of STRs (51.4%) and splicing events (70.53%) from significant spliceSTRs in HBCC could be tested (**Methods**; **Fig. S4c**). For the overlapping spliceSTRs that could be tested (n=34,828), effect sizes in the HBCC dataset are significantly correlated with effect sizes in the four replication tissues (**Fig. 1c**, Pearson’s *r*>0.7, two-tailed *P*<1×10^-100^ for all tissues). HBCC effect sizes, which are based on DLPFC, are most strongly correlated with GTEx brain frontal cortex BA9 (Pearson’s r=0.75; two-sided *P*<1×10^-100^), and least correlated with GTEx whole blood (Pearson’s r=0.63; two-sided *P*<1×10^-100^; **Fig. S4d**) when considering spliceSTRs passing at FDR<5% in either dataset. We further examined spliceSTRs for which the STR is a homopolymer repeat, which are known to be challenging to genotype, and found similarly high concordance rates (Pearson’s *r*>0.7, two-tailed *P*<1×10^-100^ for all tissues; **Fig. S5**) demonstrating similar replicability of homopolymer vs. non-homopolymer spliceSTRs. Full spliceSTR summary statistics for the GTEx analysis are available in **Supplementary Dataset 2**. **Fig. 1d-e** shows an example spliceSTR for which a dinucleotide *AC* repeat (relative to the coding strand) is associated with skipping of exon 9 of *SH3YL1* (ENST00000356150.10). The same STR tested with a similar splicing event was found as significant in three of the four replication tissues (**Fig. S6**). Although this spliceSTR was identified by linear association tests, the observed relationship between STR dosage and PSI is not strictly monotonic for higher dosages. We anticipate this is driven primarily by non-linear associations of STR length with PSI (**Fig. S7**) but may also be partially explained by lower sample sizes for higher dosages at this locus.

Finally, we compared spliceSTRs identified here to those reported by recent studies that used GTEx^39^ (49 tissues; 9537 total spliceSTRs) and multiple brain datasets (8 tissues; number of spliceSTRs range 25,033 - 270,212 across datasets considered)^40^. Similar to above, due to differences in the tools used for TR calling and splicing detection (**Fig. S8a**) and differences in the sets of spliceSTRs for which summary statistics are available from other studies (all tested pairs in GTEx vs. only significant pairs for the other brain datasets) only a small subset of spliceSTRs (range 1067-4825) that were identified in HBCC overlapped with those reported across the 57 tissues in these two studies. Our estimated effect sizes of those shared spliceSTRs are significantly correlated (Pearson’s *r*>0.47, *P*<1×10^-64^ for all tissues) with those reported by these studies (**Fig. S8b**), with the highest correlation for spinal cord lumbar (Pearson’s *r*=0.62, two-sided *P*<1×10^-100^) and similarly high correlation with frontal cortex from GTEx (*r*=0.59, two-sided *P*=7.0×10^-213^), NYGCALS (*r*=0.61, two-sided *P*<1×10^-100^) and ROSMAP (*r*=0.59, two-sided *P*<1×10^-100^). Similar results were obtained when assessing the replication of significant spliceSTRs reported by other studies in the dataset analyzed here: we identified between 26-9,242 previously reported spliceSTRs which were tested in our study, with highly concordant effect sizes reported across studies (Pearson r range 0.50-0.79; **Fig. S9**). We additionally examined the relationship between imputation quality and spliceSTR replication rates. We observed that spliceSTRs driven by STRs with higher imputation accuracy tend to have stronger concordance of effect sizes across datasets (**Fig. S10a**) and overall higher effect sizes (**Fig. S10b**). However, similar trends were observed when considering spliceSTRs estimated from GTEx for which WGS-derived STR genotypes were available (**Fig. S10b**), suggesting this trend may be driven by other factors besides imputation quality alone such as the lower polymorphism rates and stronger linkage disequilibrium (LD) with other potential causal variants for the most easily imputed STRs^43,44^.

### Fine-mapping using multiple orthogonal methods identifies putatively causal spliceSTRs

The analyses above identified 25,230 unique splicing events that were significantly associated with the lengths of nearby STRs. However, STRs are often in high LD with nearby SNPs or other variants^45^, and therefore a subset of the identified spliceSTRs could actually be driven by causal effects of different nearby variants such as SNPs that are included in the reference haplotype panel we used for imputation. To enable joint fine-mapping of STRs and SNPs, we conducted association tests between SNPs and each of these splicing events, similarly considering all available SNPs within 100kb of the two constitutive exons for each splicing event. As expected, for all but 44 of the 25,230 events for which a significant spliceSTR was identified, we also detected a significant association signal for at least one SNP (**Fig. S11a**). The 44 events with only STR signals showed overall weaker association statistics and are disregarded hereafter (**Fig. S11b**).

For the events with both significant STRs and SNPs, we next sought to identify which might be causally driven by STRs. Based on our previous observations that different fine-mappers can output inconsistent results, particularly when including STRs with lower imputation accuracy^27^, we applied three orthogonal methods in parallel for this task – analysis of variance (ANOVA), FINEMAP^46^ and Lasso regression (**Fig. 2a**; **Methods**). Consistent with our previous results on gene expression data at similar sample sizes^26^, we found that for many events no single variant was confidently fine-mapped by FINEMAP (**Fig. S12**). Despite the uncertainty from individual fine-mappers, we found that the three methods showed overall consistent results (**Fig. S13**). For example, spliceSTRs with stronger ANOVA *P*-values tended to have larger FINEMAP PIPs and were more likely to be selected by Lasso regression (**Fig. S13a-b**). Overall, we identified 17,524 spliceSTRs passing ANOVA (FDR<10%), 4,647 passing FINEMAP (PIP>0.5), and 7,830 passing Lasso (largest coefficient; **Methods**; **Fig. S14; Supplementary Dataset 1**). For downstream analysis, we defined two sets: (1) leniently fine-mapped spliceSTRs (n=21,893) that passed at least one fine-mapping method based on the criteria above and (2) strictly fine-mapped spliceSTRs (n=1,313) that passed all three methods. Within these sets, 74.8% (leniently fine-mapped) and 68.8% (strictly fine-mapped) consist of homopolymer repeats (**Fig. S15**). **Fig. 2b** shows an example of a strictly fine-mapped spliceSTR, consisting of a homopolymer *T* repeat (relative to the coding strand, hg38:chr12:69486183) that is identified as a strong potential causal variant for skipping of exon 2 of *FRS2* (ENST00000550316.5). Additional fine-mapped examples from the lenient group that pass at least two fine-mapping approaches are shown in **Fig. S16**.

**Figure 2.**
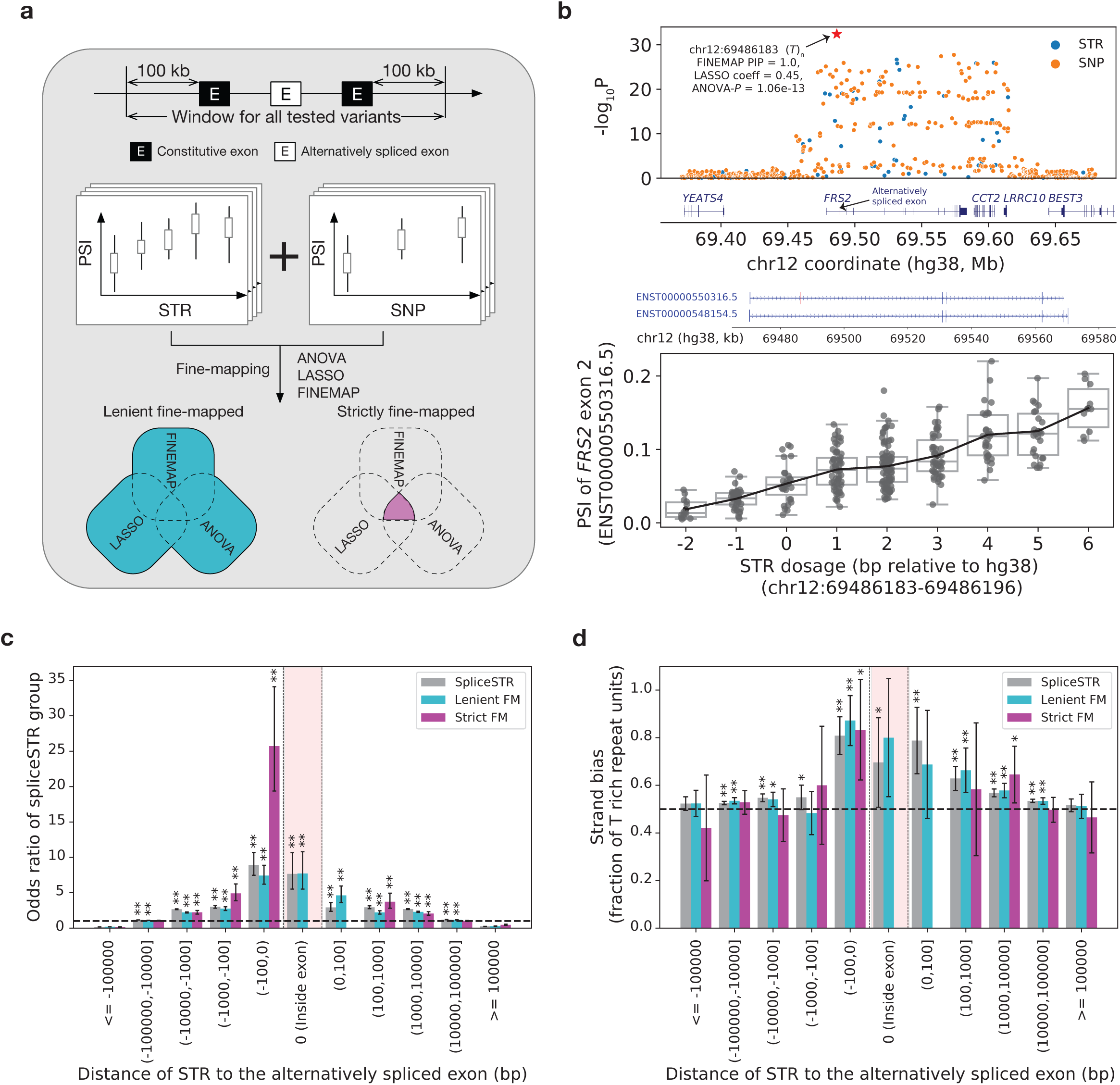
Characterization of fine-mapped spliceSTRs identified using multiple orthogonal methods. **(a) Schematic overview of the classification of leniently and strictly fine-mapped spliceSTRs.** We applied three orthogonal fine-mapping methods to prioritize potential causal variants for splicing events with significant spliceSTRs which were used to define leniently (passing at least one method) and strictly (passing all) fine-mapped spliceSTR sets. **(b) Example fine-mapped spliceSTR.** The upper panel shows association of variants with PSI of the skipped exon 2 of *FRS2* (ENST00000550316.5). The *x*-axis represents genomic coordinates on the hg38 reference. The *y*-axis represents association -log_10_ *P*-values. Blue=STRs; orange=SNPs; red star=fine-mapped STR. The bottom panel shows STR dosage (*x*-axis) vs. PSI of exon 2 of *FRS2* (ENST00000550316.5) (*y*-axis). The full exonic structure of two representative FRS2 transcripts, and the exon involved in the splice event (marked by a pink box) are shown between the two panels. **(c) Enrichment of spliceSTR categories binned by the distance to the nearest splice site of the target exon.** SpliceSTRs in each category were binned based on their stranded distance to the closest splice site of the target exon. The *y*-axis represents the odds ratio comparing each group of spliceSTRs to all tested STRs in each bin. Error bars represent ± 1 s.e. Asterisks indicate statistically significant enrichment (one-tailed Fisher’s exact test) for each spliceSTR category. **(d) Strand bias of spliceSTR categories.** The *x*-axis is the same as panel **(c)**. The *y*-axis represents the fraction of repeat units with strand bias toward “T” for each spliceSTR in each bin. Error bars represent 95% confidence intervals. Asterisks indicate the fraction of “T” repeat units significantly greater than 50% (one-tailed binomial test). For **c-d**, grey=all; cyan=leniently fine-mapped; purple=strictly fine-mapped; pink box=target exon; * indicates nominal *P*<0.05; ** indicates adjusted *P*<0.05.

Although all three fine-mapping approaches used above are designed to handle some collinearity between variants, Lasso regression and FINEMAP in particular may show instability in the case of perfect or near perfect correlation between input variants. To further evaluate the robustness of our fine-mapping results, we performed bi-directional conditional regression, in which we repeated association tests conditioning on either the fine-mapped STR or the top associated SNP for each of the n=1,313 strictly fine-mapped spliceSTRs (**Fig. S17**). We observed that all but one fine-mapped spliceSTR (n=1,312) remained nominally significant (P<0.05) after conditioning on the best SNP, compared to 63.5% (n=833) of best SNPs after conditioning on the corresponding fine-mapped STR. For many splice events, STRs show stronger associations after conditioning on the best SNP compared to SNPs conditioned on the best STR (**Fig. S17**), providing further evidence that our fine-mapping results prioritize STRs that are stronger potential candidates than other tested variants in the region. Conditional regression results for three example spliceSTRs are shown in **Fig. S18**.

We hypothesized that if spliceSTRs identified by the strategies above are truly causal, they should be enriched for certain genomic characteristics compared to non-spliceSTRs, and that enrichments should be strongest in the stringent fine-mapped set. We first compared the genomic distributions of each spliceSTR category to all tested STRs. As expected, candidate causal spliceSTRs show significant enrichment within the transcribed regions of their target genes relative to non-transcribed regions (two-tailed Fisher’s Exact Test odds ratio=2.2, *P*=5.3x10^-36^ for the strict set; odds ratio = 2.3, *P*<1x10^-200^ for the lenient set; **Fig. S19**). We further stratified all tested STRs by their distance to the alternatively spliced exon and performed the same enrichment analysis as above. We found that candidate casual spliceSTRs are most strongly enriched in regions closest to the affected exon. This trend was strongest for strictly fine-mapped spliceSTRs within 100 base pairs upstream of the affected exons (two-tailed Fisher’s Exact Test odds ratio=25.9, *P*=2.8×10^-15^ vs. odds ratio=7.3, *P*=1.4×10^-20^ for the leniently fine-mapped spliceSTRs, **Fig. 2c**). Pathway analysis did not identify any enriched pathways in genes containing fine-mapped spliceSTRs vs. all tested genes (**Methods**) but identified enrichment of “pathways of neurodegeneration-multiple diseases” (FDR 1.8%) and “metabolic pathways” (FDR 1.8%) when comparing genes with any significant spliceSTR vs. all tested genes.

Next, we asked whether fine-mapped spliceSTRs have different sequence characteristics compared to other tested STRs. We examined the total length (in number of repeat copies) based on the hg38 reference sequence and found that fine-mapped spliceSTRs have significantly higher copy numbers compared to spliceSTRs not passing fine-mapping (Mann-Whitney adjusted one-sided *P*<0.05 for both the strict and lenient sets; **Fig. S20a**). When stratifying STRs by repeat unit length, this trend is most significant for homopolymers but remains nominally significant for most of the other repeat unit lengths (**Fig. S20b**), which is likely due to reduced power arising from the relatively small number of spliceSTRs with longer repeat units. The magnitude of the difference in length across all units is strongest for the strictly fine-mapped set (**Fig. S20**). The tendency of fine-mapped STRs to be longer may be driven in part by the fact that longer STRs tend to exhibit higher polymorphism rates in the population resulting in weaker LD with SNPs^44^ and therefore making them easier to fine-map. Indeed, among all STRs significantly associated with splicing events, those with greater heterozygosity showed greater overlap with candidate causal spliceSTRs defined by both fine-mapped sets (**Fig. S21**).

We further assessed whether fine-mapped spliceSTRs are enriched for individual repeat unit sequences (see **Methods**). In general, we found *CG*-rich repeats (e.g., *CCG* with odds ratio=10.6 and *CGG* with odds ratio=5.7) were significantly enriched (two-tailed Fisher exact test *P*<0.05) in the strictly fine-mapped sets compared to all STRs tested. Overall, enrichments were directionally consistent between the two fine-mapped sets, with higher odds ratios observed compared to all significant spliceSTRs. Many repeat units showed suggestive enrichment but did not reach statistical significance, which we hypothesize in many cases is because of low counts (**Fig. S22**). Next, given previous observations of a strand bias toward “T”-rich (vs. “A”-rich) units on the coding strand within transcribed regions^47^, we additionally tested whether spliceSTRs show a similar strand bias. We found a consistent bias toward “T” rich orientations of repeat units relative to the transcribed strand of associated genes across all sets of STRs tested regardless of fine-mapping status (**Fig. 2d**), suggesting evolutionary pressure constraining sequence content in this region. This bias is strongest for STRs located within 100 base pairs of the affected exon (one-sided binomial *P*=1.1×10^-6^ for the leniently fine-mapped set and *P*=0.02 for the strictly fine-mapped), which is the same region where strictly fine-mapped STRs are most enriched.

Taken together, our results identify distinct characteristics of candidate causal spliceSTRs, including a subset that are primarily long, T-rich sequences in close proximity upstream of target exons. Our observations that many of these characteristics are most strongly enriched in strictly fine-mapped spliceSTRs suggests that our fine-mapping strategies are indeed enriching for STRs with causal effects on splicing.

### Fine-mapped spliceSTRs are enriched near binding sites of RNA-binding proteins (RBPs)

Previous studies have shown that STRs are the most frequently bound class of repetitive elements by RNA binding proteins (RBPs)^39,48^, and that differences in STR copy number may impact RBP binding and alter RBP-mediated splicing^49–53^. To evaluate whether RBP binding is a potential mechanism driving the observed spliceSTRs, we first assessed the overlap between STRs in our dataset with RBP binding sites using eCLIP data available for 129 distinct RBPs from both K562 and HepG2 cell lines^48^, restricting to RBPs that are located on autosomes and expressed in the HBCC DLPFC dataset (**Methods**). All STRs overlapping RBP binding peaks were stratified by repeat unit, followed by visualization of the eCLIP read coverage of the associated RBPs. We observed that STRs are often located either within the RBP binding sites or in the flanking region directly adjacent (**Fig. S23**), similar to findings from previous work^54^, but show distinct patterns for different RBP-repeat unit pairs. In some cases, particularly homopolymer (*T)*_n_ repeats, we found strong strand preferences in which RBP coverage is strongest downstream of the STR, although this strand preference could in some cases be driven by biases in the eCLIP protocol (**Discussion**).

We observed that leniently fine-mapped spliceSTRs are significantly enriched (two-tailed Fisher’s Exact Test adjusted *P*<0.05) for overlapping binding sites for 32 RBPs, with fewer significantly enriched RBPs (n=6) obtained from the strictly fine-mapped spliceSTRs, likely due to the lower counts of that set (**Fig. 3a**). These observations are consistent with a previous spliceSTR study which identified an enrichment for spliceSTRs overlapping RBP binding sites^39^. Using previously published data from Boyle et al.^48^, we annotated RBPs according to the typical genomic locations of their binding sites and found that the majority of enriched RBPs are those that tend to bind in intronic regions and coding sequences (CDS), likely because of the larger number of RBPs in these two categories. Compared to all tested STRs, enrichments for spliceSTRs overlapping RBP binding sites are overall higher when we restrict analysis to STRs that are located within the constitutive exons of the splicing events (**Fig. S24**), which is in line with the finding that the impact of RBPs on splicing decays as their distance to the splice site increases^55^. We further categorized RBPs based on whether they are involved in splicing regulation or form components of the spliceosome, based on previously reported annotations^56^ (**Fig. 3a**). Compared with all tested STRs, both strictly and leniently fine-mapped spliceSTRs were significantly enriched for overlapping binding sites of spliceosomal components (strict FM spliceSTRs: two-tailed Fisher’s exact test odds ratio=3.54, *P*=1.8×10^-8^; lenient FM spliceSTRs: odds ratio=2.20, *P*=1.3×10^-25^) and to a lesser extent for RBPs involved in splicing regulation (strict FM spliceSTRs: odds ratio=1.76, P=1.1×10^-5^; lenient FM spliceSTRs: odds ratio=1.72, *P*=3.4×10^-50^).

**Figure 3.**
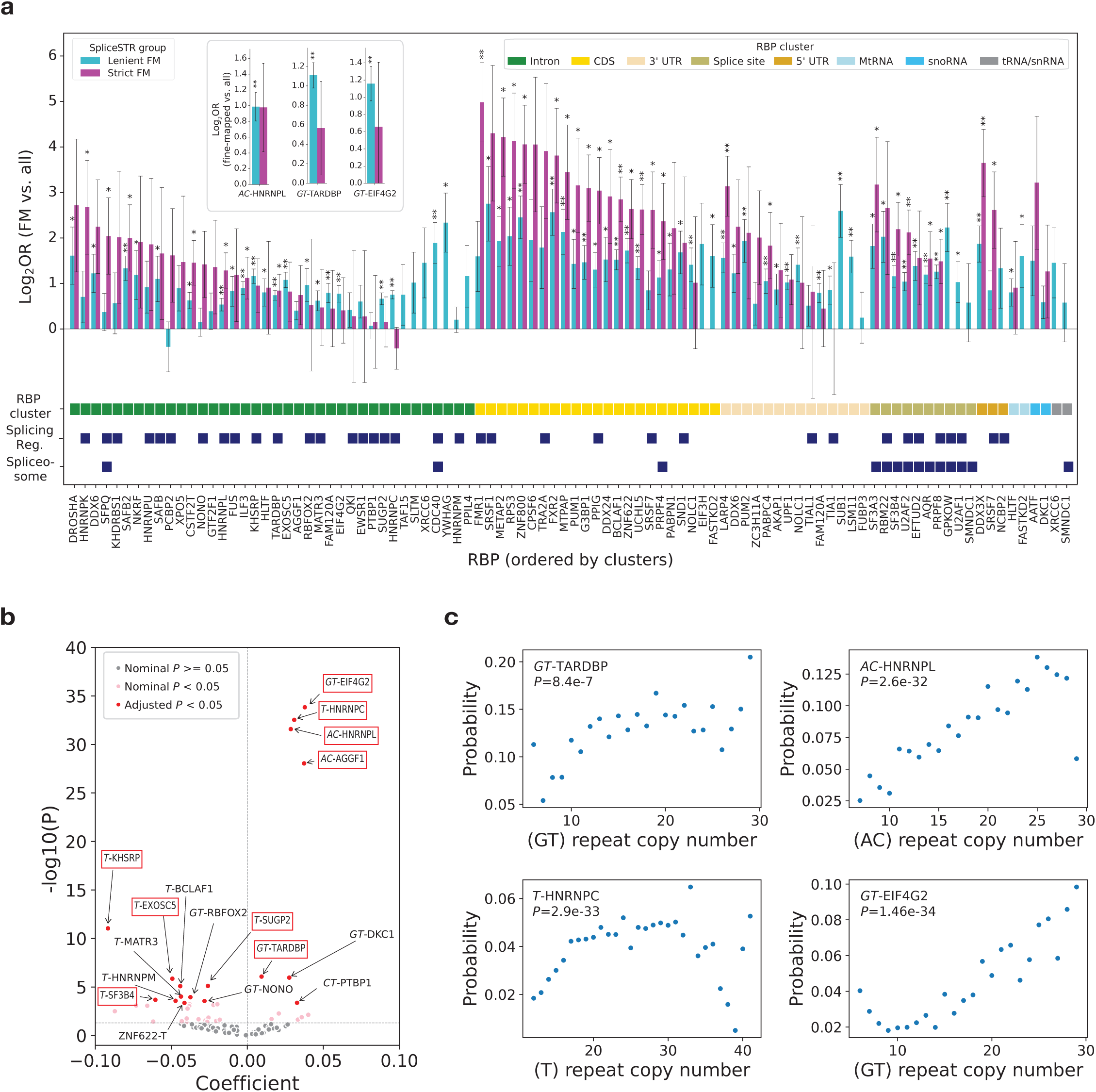
Characterizing enrichment of fine-mapped spliceSTRs at RBP binding sites. **(a) Enrichment of RBP binding sites among fine-mapped spliceSTRs.** The *x*-axis represents all RBPs tested for which binding sites (Boyle *et al.*^48^) overlapped at least 3 unique leniently fine-mapped spliceSTRs. The *y*-axis denotes the log_2_ odds ratio comparing each fine-mapped spliceSTR set to all tested STRs for each RBP. For the inset panel, the *x*-axis represents stranded repeat units and the *y*-axis similarly denotes the log_2_ odds ratio. Cyan=leniently fine-mapped spliceSTRs, purple=strictly fine-mapped spliceSTRs. Error bars represent ± 1 s.e. Asterisks indicate statistically significant enrichment or depletion (two-tailed Fisher’s exact test, * nominal *P*<0.05, ** adjusted *P*<0.05) for each group. RBPs are grouped according to annotated genomic binding location patterns^48^, with groups indicated by different colors as shown in the RBP cluster legend. RBPs that are spliceosomal components or involved in splicing regulation (from Rhine *et al.*^60^) are highlighted with dark blue boxes. **(b) Association between repeat unit copy number and RBP binding probability.** Each point represents a repeat unit-RBP pair, with colors indicating the -log10 *P*-values (obtained by logistic regression). Repeat units are stranded, based on the sequence relative to the coding strand. The *x*-axis shows coefficients from logistic regression. The *y*-axis indicates the -log_10_(*P*-values). Significantly associated repeat unit–RBP pairs (adjusted P < 0.05) are labeled, with red boxes indicating pairs that are also significantly enriched in leniently fine-mapped vs. all spliceSTRs. **(c) Examples of significantly associated RBP-repeat unit pairs.** The *x*-axis represents the repeat unit copy number according to the reference genome. The *y*-axis shows the probability of RBP binding for STRs in each bin. Nominal *P*-values from the logistic regression testing the association between repeat unit length (in copy number based on hg38 coordinates) and the probability that the repeat overlaps with RBP binding sites are labeled in each plot.

To test whether enrichments for certain RBPs are associated with STRs with specific sequences, we further stratified all tested STRs by their repeat unit and performed the same enrichment analysis as above. While no significant repeat unit–RBP pairs were detected in the strictly fine-mapped set, likely due to limited statistical power, we identified a total of 15 repeat unit-RBP pairs (corresponding to 5 unique repeat units and 13 unique RBPs) for which leniently fine-mapped spliceSTRs are significantly enriched (two-tailed Fisher’s Exact Test adjusted *P*<0.05) for overlapping RBP binding sites compared to other tested STRs. Among these pairs, (*T)*_n_ is the most common, showing significant enrichment with 7 different RBPs. Of the 15 repeat unit RBP pairs identified here, 9 were previously reported^48,57^ including the well-known *GT*-binding factor TARDBP (two-tailed Fisher’s exact test odds ratio=2.2, adjusted *P*=4.0×10^-13^), the *AC*-binding factor HNRNPL (two-tailed Fisher’s exact test odds ratio=2.0, adjusted *P*=7.0×10^-5^) and the *GT*-binding factor EIF4G2 (two-tailed Fisher’s exact test odds ratio=2.2, adjusted *P*=1.8×10^-5^) (**Fig. 3a; Fig. S25**).

The analyses above were performed using RBP binding sites obtained from K562 and HepG2 whereas the spliceSTRs are based on the dorsolateral prefrontal cortex. While prior studies have shown that many RBPs exhibit conserved sequence motifs and binding preferences across cell types^58,59^, we sought to assess this using eCLIP datasets available for a small number of RBPs for more brain-relevant cell types. For this, we analyzed eCLIP data for two RBPs: G3BP1 from human frontal cortex and human induced pluripotent stem cell (hiPSC)- derived neurons, and TARDBP from hiPSC-derived neurons^60^. We observed similar enrichment patterns of fine-mapped spliceSTRs overlapping RBP binding sites compared to those identified in the cell line data (**Fig. S26**).

Although the analyses above demonstrate enrichment for spliceSTRs within RBP binding sites, they do not directly test the impact of STR copy number on RBP binding. To assess whether variation in repeat copy number might impact RBP binding in the HepG2 and K562 datasets, we first tested for a genome-wide relationship between the total length of each repeat (based on the hg38 coordinates) and the fraction of STRs of that length overlapping an RBP binding site, stratified by repeat unit sequence (**Methods**). Overall, we found that 17 out of 98 tested repeat unit-RBP pairs showed significant association (adjusted *P*<0.05) (**Methods; Fig. 3b**). Of these, 7/17 showed positive associations, in which longer repeats are more likely to be bound by the target RBP. For example, we found that HNRNPL binding is positively associated with the length of (*AC*)_n_ repeats (adjusted *P*=3.1×10^-30^), consistent with published results based on RNA affinity purification and pull-down assay^52,61^. We also found that the length of (*GT)*_n_ repeats is positively correlated with the probability of TARDBP binding (**Fig. 3c**; adjusted *P*=9.9×10^-5^ based on logistic regression; **Methods**), again matching expectations based on previous literature^62–64^. This trend is recapitulated in the hiPSC-derived neuron eCLIP dataset for TARDBP (*P*=0.04). Although not initially expected, we observed many cases in which repeat length is negatively associated with RBP binding probability (**Fig. S27**). One possible explanation is that such negative correlations may be partially driven by challenges in uniquely mapping short eCLIP reads to regions with longer repeat tracts, which could lead to an underestimation of RBP binding signals at longer repeats. It is also possible that positive correlations could be related to other types of mapping artifacts at repeat regions, such as incorrect mapping of highly repetitive reads. Additional examples of both positive and negative associations are shown in **Fig. S27**. The set of repeat unit-RBP pairs for which repeat length is associated with RBP binding shows significant overlap with repeat unit-RBP pairs enriched in leniently fine-mapped vs. all spliceSTRs (two-tailed Fisher’s exact test odds ratio=11.1, *P*=2.6×10^-4^), although this enrichment may be partially driven by power as repeat unit-RBP pairs with more binding sites tend to be identified in both association and enrichment analysis. Taken together, our results suggest a subset of spliceSTRs may be driven by STRs with specific repeat units that modulate binding of RBPs as a function of STR length.

### Characterizing spliceSTRs underlying GWAS signals for brain-related complex traits

To determine whether fine-mapped splicing events might underlie GWAS signals for brain-related traits, we performed colocalization analysis using the fine-mapped splicing events and published GWAS summary statistics for six disorders: schizophrenia (SCZ), autism spectrum disorders (ASD), bipolar disorder (BD), attention deficit/hyperactivity disorder (ADHD), major depressive disorder (MDD) and Alzheimer’s disease (AD; **Fig. 4a**). We considered all splicing events with a leniently fine-mapped spliceSTR located within 1 megabase (Mb) of lead variants for published GWAS signals for these traits. In total, we tested 6,111 splicing event-trait pairs (**Supplementary Table 1**). We identified candidate co-localized pairs as those with colocalization posterior probability for a shared causal variant (PP.H4)>50%. Because our colocalization analysis does not directly consider STRs, since those do not have GWAS summary statistics available, we additionally restricted to pairs for which the fine-mapped spliceSTR is in at least moderate LD (*r^2^*>0.1) with the lead GWAS variant in the region for that trait. As many fine-mapped splice events map to the same gene, we aggregate colocalized fine-mapped splice events at the gene level to avoid counting events from the same gene multiple times. In total, we identified 96 gene-trait pairs passing these criteria (**Fig. 4b**), containing 11 strictly fine-mapped pairs. Schizophrenia showed the highest number of colocalized genes (60 splice events; PP.H4>50%), although the high number of colocalized splicing events is likely driven by the higher total number of GWAS signals and GWAS sample sizes for that trait. As expected, the total number of such co-localized events identified varies with the PPH4 and LD thresholds used, although this trend is less pronounced for the strictly fine-mapped set (**Fig. S28**).

**Figure 4.**
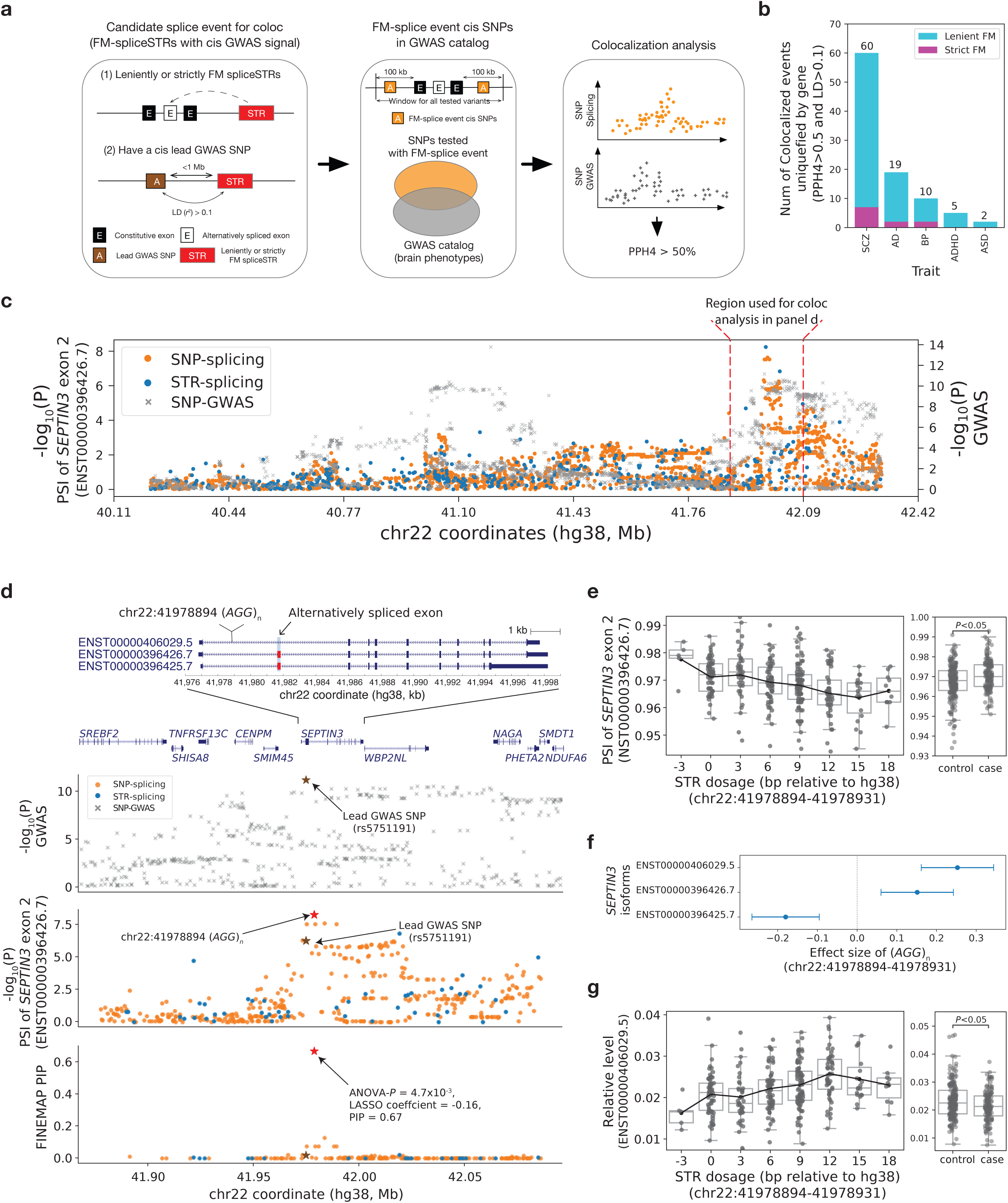
Colocalization of fine-mapped spliceSTRs with GWAS signals. **(a) Overview of colocalization workflow.** We identified splice events with PP.H4>0.5 based on SNP splicing and GWAS summary statistics for which a fine-mapped spliceSTR is in LD (*r^2^*>0.1) with the lead GWAS variant. **(b) Gene-level counts of fine-mapped colocalized splice events.** Bars show the number of genes with colocalized fine-mapped splice events for each trait (purple=strictly; cyan=leniently fine-mapped spliceSTRs). MDD not shown since no colocalized events were identified. **(c) Summary statistics for *SEPTIN3* exon 2 PSI and schizophrenia.** The left *y*-axis shows -log_10_(*P*-values) for skipping of *SEPTIN3* exon 2 (ENST00000396426.7) (orange=SNPs; blue=STRs). The right *y*-axis shows -log_10_(*P*-values) for schizophrenia^70^. Red dashed lines indicate the window used for splicing association tests. The *x*-axis shows hg38 coordinates. **(d) Summary statistics for the *SEPTIN3* region.** Panels, from top to bottom, provide genomic annotations, -log_10_(*P*-values) for schizophrenia, -log_10_(*P*-values) for splicing, and FINEMAP PIPs for all nominally significant SNPs and STRs associated with exon 2 skipping for the highlighted window in **c**. All transcripts whose expression is significantly associated with the (*AGG*)n STR (hg38:chr22:41978894-41978931) are shown. **(e) Association between STR dosage and PSI.** Left panel: the *x*-axis represents dosage of the (*AGG*)n STR. The *y*-axis represents PSI of exon 2. Each dot represents an individual. Box plots summarize distributions of PSI values. Boxplot elements are same as in Fig. 1d. Black line shows mean expression for each *x*-axis value. Right panel: PSI of exon 2 in controls (n=194) vs. cases (BD or schizophrenia, n=142). **(f) *SEPTIN3* isoform associations.** The plot shows *SEPTIN3* isoforms significantly associated (nominal *P*<0.05) with the (*AGG*)n STR. The *x*-axis indicates effect size and 95% confidence intervals. **(g) Association of STR dosage vs. *SEPTIN3* isoform ENST00000406029.5.** The left panel shows the relative abundance of transcript ENST00000406029.5 vs. STR dosage. The right panel compares relative transcript abundance in cases vs. controls.

Our colocalized fine-mapped spliceSTRs recapitulate previously reported signals, supporting the validity of our analysis. For instance, we confirmed strong co-localization between skipping of exon 12 of *PLEKHA1* (ENST00000368989.6) with an Alzheimer’s disease GWAS signal in this region (PP.H4=88.97%)^39,40^. This splice event was strictly fine-mapped to a (*GT)*_n_ STR at hg38:chr10:122429495 with a FINEMAP PIP of 0.96 and the STR is in moderate LD (*r²*=0.71 in Europeans) with the lead GWAS SNP (rs7908662). We found this repeat is also significantly associated with the relative abundance of two isoforms of *PLEKHA1* (**Fig. S29**), suggesting that the colocalized spliceSTR may contribute to Alzheimer’s disease risk by modulating the expression of *PLEKHA1* isoforms. We additionally identified a previously reported co-localization between skipping of exon 6 of *DGKZ* (ENST00000456247.6) and schizophrenia^65^ for which we identified multiple candidate leniently fine-mapped spliceSTRs (**Fig. S30**).

Our analysis also uncovered examples of co-localization between splicing events and complex traits that, to our knowledge, have not been previously reported. Several of these involved *SEPTIN3*, a member of a highly conserved family of GTP-binding proteins implicated in synaptic dysfunction which is associated with multiple psychiatric and neurodegenerative disorders including schizophrenia, Parkinson’s disease, and Alzheimer’s disease^66–69^. We identified three separate candidate splicing signals in *SEPTIN3* that show strong colocalization with schizophrenia (GWAS signal lead SNP rs5751191). The first and strongest involves skipping of exon 2 of *SEPTIN3* (ENST00000396426.7); colocalization PP.H4=65.3%; **Fig. 4c**), which was strictly fine-mapped by all three methods to an intronic (*AGG*)_n_ (relative to coding strand) STR at hg38:chr22:41978894 (FINEMAP PIP=0.67, LASSO coefficient=-0.16, ANOVA *P*-values=4.7×10^-3^) that is 2.7kb upstream of the affected exon. This repeat is negatively correlated with inclusion of exon 2 and positively correlated with the relative abundance of isoform ENST00000406029.5 which excludes that exon (**Fig. 4e-f**). We also observe opposite associations of the same repeat with transcripts ENST00000396426.7 and ENST00000396425.7, despite both containing exon 2. This likely reflects the fact that isoform formation arises from combinations of multiple splicing events (**Fig. S31**), such that the same exon can be incorporated into distinct transcript contexts with different regulatory outcomes. Notably, we found that the alternative splicing of exon 2 in this event differs significantly between cases and controls in HBCC (n=60/82 bipolar/schizophrenia cases, n=194 controls; Mann-Whitney two-sided nominal *P*=1.3×10^-3^). The relative expression level of the associated transcript, ENST00000406029.5, also shows a significant difference between case and control groups (*P*=0.02), suggesting the skipping of exon 2 of *SEPTIN3* may be related to psychiatric disease.

Inspection of this region revealed that the GWAS signal spanned beyond the co-localization window initially considered and appears to contain multiple peaks. To account for the difference in signal window sizes between splicing and GWAS data, we extended the splicing association testing range to match the region reported based on LD clumping of the GWAS signal in this region^70^ and found that the (*AGG*)_n_ spliceSTR overlaps with one of the GWAS peaks in this region (**Fig. 4d**). Additional co-localization signals identified in this region include a mutually exclusive exon event associated with a *GT/AG* repeat in the 3’UTR of *SEPTIN3* (PPH4=94.49%) that has been previously linked to Alzheimer’s disease^66^ and a separate skipping event of exons 2 and 3 that is expected to induce a frameshift and early stop^66^ associated with a homopolymer repeat (PPH4=89.46%). Although evidence for co-localization of these second two splicing events with schizophrenia is strong, fine-mapping evidence for their corresponding STRs are weaker than for the *AGG* repeat (both pass ANOVA but not FINEMAP or Lasso). Additional examples of co-localization between fine-mapped splice events and complex traits are shown in **Fig. S32**. Overall, these findings highlight the potential contribution of STRs to the complex splicing regulation of *SEPTIN3* and its role in disease pathogenesis.

The analysis above relies on colocalization analysis, in which we indirectly identify spliceSTRs that may underly GWAS signals based on LD patterns with nearby SNPs. To enhance our ability to directly detect disease-associated spliceSTRs, we examined whether STRs recently identified as associated with different diseases overlapped our spliceSTR set. Notably, although multiple studies have shown STRs have been implicated in both schizophrenia and autism, these studies have focused on rare repeat expansions^71–73^ and *de novo* variants^73,74^ that are not represented in our callset which focuses on common inherited variants. While we identified 63 spliceSTRs (21 unique STRs) overlapping STRs previously tested for expansions in autism^73^, none showed significant outlier expansions in those cohorts. We next compared our spliceSTR catalog with a recently published genome-wide STR association study in Alzheimer’s Disease (AD)^75^. Among STRs tested in both studies, we identified 19 AD-associated STRs (P<5e-8) that are also significant spliceSTRs, most of which fall on Chromosome 19 near the APOE region (**Supplementary Table 2**). These include a homopolymer G repeat at hg38:chr19:44909968 that is leniently fine-mapped for *TOMM40* splicing in our analysis and strongly associated with AD (*P*=7.65×10^-1201^). In summary, while we currently find limited overlap of spliceSTRs with STRs that have been directly associated with disease, we hypothesize that this is largely due to the small number of STR association studies to date and anticipate the overlap will grow as additional genome-wide STR studies are performed.

## Discussion

In this study, we leveraged deep RNA-seq data from dorsolateral prefrontal cortex (DLPFC) and imputed STR genotypes from 336 individuals to generate a detailed resource of 146,969 splicing-associated short tandem repeats (spliceSTRs). We performed fine-mapping using three orthogonal methods to prioritize potential causal spliceSTRs and characterized the top candidates. Finally, we identified examples of fine-mapped spliceSTRs with strong evidence of co-localization with GWAS signals for brain-related disorders, suggesting they play a role in driving complex trait risk.

We found fine-mapped spliceSTRs were strongly enriched within 100 bp upstream of their affected exons, suggesting these spliceSTRs may influence splicing through local *cis*-regulatory mechanisms. This region frequently overlaps, or lies adjacent to the polypyrimidine tract, a *U/C*-rich sequence that is critical for spliceosome assembly^76,77^. The observed bias toward “*T*” rich repeat units on the transcribed strand is therefore consistent with functional constraints on polypyrimidine tract composition. Variation in STR length at these loci could modulate the effective length or sequence composition of the polypyrimidine tract, altering splice factor binding affinity or splice site recognition and ultimately impacting exon inclusion.

Consistent with previous reports of the widespread binding of RBPs to repetitive sequences^48,52,58,78,79^, fine-mapped spliceSTRs were significantly enriched for overlap with RBP binding sites compared to non–fine-mapped spliceSTRs. Notably, several of the enriched RBPs, such as TARDBP^80,81^, are well known for their roles in splicing regulation in the human brain, while the functions of others remain less understood. Genome-wide analysis of RBP binding sites showed unique positional patterns of RBP bindings for specific RBP-repeat unit pairs (**Fig. S23**), particularly for homopolymer *(T)_n_* repeats. Onoguchi *et al*.^54^ observed similar biases and showed potential stable secondary structures are often located upstream of eCLIP peak summits. However, given known positional biases introduced by eCLIP library preparation, such as RNase digestion and reverse transcription truncation^82–84^, we note that observed downstream shifts in peak coverage may be driven in part by those experimental artifacts. We additionally identified multiple strong relationships between RBP binding and repeat length, including known associations between HNRNP-L and AC repeats^52^, between TARDBP and GT repeats^80^, and additional examples many of which have not been previously reported (**Fig. 3b**). Further validation of these trends across a broader set of RBPs in human brain tissue will be important as such datasets become available. Additionally, altered RBP binding at different repeat lengths could reflect a combination of effects driven by repeat length, sequence composition of imperfect repeats, or effects on RNA secondary structure and stability, and we are not able to explicitly distinguish between these different factors here. Although our results suggest a subset of spliceSTRs act by modulating binding affinity of RBPs, by which they impact splicing efficiency of nearby exons, not all spliceSTRs are explained by this mechanism. Additional mechanisms, such as effects mediated by RNA or DNA folding energy and the formation of DNA or RNA secondary structures, remain to be explored on a genome-wide scale.

Although the HBCC includes schizophrenia and bipolar disorder cases, sample sizes are insufficient to directly perform GWAS in this dataset. Instead, by leveraging publicly available GWAS summary statistics for brain-related diseases, we found evidence that a subset of fine-mapped spliceSTRs show strong co-localization with disease-associated GWAS loci. Several of these, including a spliceSTR for *PLEKHA1* co-localized with Alzheimer’s disease, were previously identified^40^. We also identified a confidently fine-mapped spliceSTR for which an intronic *(AGG)_n_*repeat is associated with PSI of exon 2 in *SEPTIN3* that is co-localized with a schizophrenia GWAS signal. *SEPTIN3* and other septins play a role in axonal transport, vesicular trafficking, and neurotransmitter release^68^. Changes in septin expression have been previously linked to both bipolar disorder^85^ and schizophrenia^67^, making this a plausible candidate gene. The repeat does not overlap RBP binding sites analyzed here. However, AGG repeats have been shown to form G-quadruplex structures^86^, which can influence splicing^87^, suggesting formation of non-canonical secondary structures as a potential mechanism. Importantly, all of these co-localization events were identified indirectly by leveraging SNP-STR LD and available SNP summary statistics. Future work is needed to directly genotype these and other STRs in large cohorts to verify these associations and determine if STRs are causally driving these signals.

Our study complements other recent genome-wide investigations of the role of STRs in splicing^39,40^, while also highlighting challenges in comparing spliceSTRs across studies. Here, we used STR genotypes imputed from a reference panel based on genotypes obtained using HipSTR^19^, which considers STRs with repeat units 1-6bp and includes repeats that have sequence imperfections or those with multiple different repeat units. We also quantified splicing events from deep DLPFC RNA-seq data using rMATs^42^, which uses exon-based detection of splicing events. In contrast, Hamanaka *et al.*^39^ used GangSTR^88^ for STR genotyping, which considers only perfect repeats, and analyzed data from GTEx which has low sample sizes for brain tissues. Cui *et al.*^40^ focused on brain tissues but excluded homopolymer STRs which comprise around 50% of the HipSTR reference. Both other studies used alternative intron-based methods for detecting splicing events. Further, differences in available genotype or WGS data, RNA-seq coverage, and read lengths resulted in differential detection of splicing events across datasets even when using the same pipeline (**Fig. S4**). Only a small fraction of spliceSTRs identified could therefore be directly compared across studies (**Fig. S8-9**). As a result, the majority of the 146,969 significant spliceSTRs detected here represent novel associations.

Our study has several limitations. First, while we analyzed hundreds of deep RNA-seq datasets for DLPFC, the sample size was relatively small and derived from a single brain region. Because many splicing events are tissue-specific, our analysis may have missed splicing variation occurring in other tissues. Further, we have limited ability to detect rare splicing events. Additionally, the cohort primarily consists of individuals of European ancestry, which may limit the generalizability of our findings to other populations. Second, our analysis relied on imputed genotypes using a reference haplotype panel based on HipSTR calls^43^ which is limited to capturing repeats that can be spanned by short read lengths. Other classes of tandem repeats (TRs), such as large repeat expansions and variable-number tandem repeats, were not included. Our previous validation of this panel shows high concordance (97% across all loci) between imputed and directly genotyped STR genotypes^27,43^. However, imputation accuracy varies considerably across loci, with highly polymorphic loci showing lower overall concordance. Ultimately, specific loci of interest may require confirmation in orthogonal datasets for which STRs can be directly genotyped. Future studies may overcome this limitation by genotyping STRs or other repeat types directly from genome sequencing data when available. Third, although we identify strong evidence that a subset of STRs are driven by interaction with RBPs, future experimental validation will be necessary to confirm this and other potential mechanisms by which STR variation influences splicing. Fourth, our colocalization analyses identify fine-mapped splicing events that colocalize with GWAS signals via SNP-based splice QTL and GWAS followed by LD mapping to STRs, which may vary across populations. Therefore, additional validation is required to confirm the contribution of colocalized fine-mapped STRs to the observed GWAS signals.

### Conclusions

Altogether, our study provides a valuable resource for studying the role of STRs in splicing. We identify thousands of spliceSTRs and demonstrate their potential role in driving risk for brain-related traits. Our results suggest that integrating STRs into genome-wide studies of variation in splicing and other molecular phenotypes is likely to uncover a rich source of loci contributing to complex traits.

## Methods

### STR imputation and genotyping

We downloaded hard-called SNPs and short indel genotypes for 364 samples that also have RNA-seq data available from the Human Brain Collection Core (HBCC) through dbGaP (phs000979.v3.p2). We used our previously published SNP-STR haplotype panel^43^ from the 1000 Genomes Project^89^ as a reference for STR imputation. We used Beagle^90^ 5.2 (version: 28Jun21.220) with the tool’s provided human genetic maps and non-default flag ‘--ap=true’ to impute STRs for each sample separately for each chromosome. For each sample, we extracted only the STR records and concatenated the VCF for each chromosome using bcftools^91^ v1.10.2. We then merged VCFs across samples using mergeSTR from TRtools v5.0.2^92^ and added back the STR INFO fields present in the SNP-STR reference panel that Beagle removed during imputation using a custom script. We lifted the coordinates from hg19 to hg38 using LiftOver^93^ with a chain file downloaded from the UCSC Genome Browser^94^. A total of 445,720 STRs were imputed and successfully lifted to hg38 coordinates.

### RNA-seq alignment and gene/isoform quantification

We downloaded the FASTQ files containing RNA-seq reads that passed QC for 364 samples from the Human Brain Collection Core (HBCC) through the Database of Genotypes and Phenotypes (dbGaP phs000979.v3.p2). Alignment and gene/isoform quantification were performed following the ENCODE RNA-seq pipeline. RNA-seq reads were aligned to the GRCh38.p13 reference genome using STAR^95^ version 2.7.9a, with annotations from GENCODE v38, to generate BAM files. Gene and isoform TPM values were quantified using RSEM^96^ v1.3.3. Genes with TPM<1 in more than 90% of samples or overlapping with segmental duplication regions obtained from the UCSC Genome Browser^94^ were removed. Analysis was restricted to protein coding genes based on GENCODE v38 annotations. For isoforms, we excluded transcripts that either had no expression or were located in segmental duplication regions. We then calculated the ratio of each isoform’s TPM to the total TPM of all isoforms for that gene.

### Quantifying alternative splicing events

We ran rMATS v4.1.2^42^ on BAM files with aligned RNA-seq reads from all 364 samples jointly to obtain exon inclusion levels (percent spliced in [PSI] values) for different splicing events based on the GENCODE v38 annotation. The relative ratio of different splicing types is comparable to these reported by rMATs^42^. We filtered three samples for which more than 20% of splicing events were not detected and samples where the imputed STR genotypes were not available due to missing SNP genotypes. The remaining 336 samples were used for downstream analysis. To ensure the reliability of estimated PSI and downstream analysis, we only included splicing events passing the following filters: 1) PSI is 0 or 100 in less than 90% of samples; 2) splicing events are supported by at least 10 total read count in more than 80% of samples; 3) the affected exon is not overlapping with any segmental duplication regions.

### Identification of SNPs and STRs associated with splicing events

For each STR within 100 kb of the two constitutive exons for a splicing event, we performed a linear regression between STR dosages and exon PSI values *Y* ∼ *β*_0_*X* + *β*_1_*C* + *ε* where X denotes STR dosages, Y denotes PSI values, C denotes covariates, *β*_0_ denotes effect size, *β*_1_ denotes coefficients for covariates, and *ε* is the error term. For each STR, we calculated dosage for each sample as the sum of the differences between the lengths of the best-guess imputed alleles and the reference alleles on both chromosomes. For loci with interrupted or mixed motifs, the estimated repeat length reflects the total length of the repeat region rather than requiring perfect motif continuity. Samples with missing genotypes or expression values were removed from each regression analysis. To reduce the effect of outlier STR genotypes, we removed samples with genotypes observed in fewer than three samples. If, after filtering samples, there were fewer than three unique genotypes, the STR was excluded from analysis. PSI values, STR genotypes and covariates for remaining samples were then Z-scaled before performing each regression to force resulting effect sizes to be between −1 and 1.

We included age, sex, RNA-seq read counts, genotype principal components (PCs), and PEER factors as covariates. To obtain genotype PCs, we performed PCA on SNP genotypes of the 423 samples from HBCC and used the top ten PCs as covariates. Analysis was restricted to biallelic SNPs and SNPs with minor allele frequency at least 5%. PCA on resulting SNP genotypes was performed using smartpca^97,98^ v.13050. PEER factors were calculated using PEER^99^ v1.0 from the z-scaled filtered PSI values with mean 0 and variance 1. We use the top 20 PEER factors as covariates.

Linear regressions were performed using the OLS function from the Python statsmodels.api v.0.14.0 module, which returns estimated regression coefficients computed using ordinary least squares and two-tailed *P* values for each regression coefficient testing the null hypothesis *β*_0_ = 0 computed from t-statistics for each coefficient. As a control, for each STR-splice event pair, we performed a permutation analysis in which sample identifiers were shuffled. We grouped STR-splice events by splicing types and adjusted *P* values using the fdrcorrection function in the statsmodels.stats.multitest module for each splicing type with an FDR threshold of 5%.

Using the same model covariates and normalization methods, we tested the association between SNPs dosages (0, 1, or 2) and PSI. To reduce computational cost, we limited SNP association testing to splicing events that were significantly associated with a STR.

### Comparison of significant spliceSTRs with GTEx tissues and other studies

To compare spliceSTRs identified in HBCC with GTEx, we obtained 30x Illumina WGS data from 652 unrelated participants from GTEx^39^ through dbGaP accession number phs000424.v8.p2. WGS data was accessed using fusera through Amazon Web Services. We genotyped autosomal STRs jointly across all 652 samples using HipSTR^19^ v0.7 with default parameters and the hg38 STR reference panel. The resulting VCFs were filtered using dumpSTR from TRTools v5.0.2^92^, using the parameters --hipstr-min-call-Q 0.9, --hipstr-min-call-DP 10, -- hipstr-max-call-DP 1000, --hipstr-max-call-flank-indel 0.15, --hipstr-max-call-stutter 0.15, --min-locus-callrate 0.5, --min-locus-hwep 0.00001, --filter-regions hg38_segDups_sorted.bed.gz (a reference of segmental duplication regions available at UCSC Genome Browser h38.genomicSuperDups table). Altogether, 990,892 autosomal STRs remained for association tests. We then downloaded the aligned RNA-seq reads for brain frontal cortex BA9, brain nucleus accumbens basal ganglia, EBV transformed lymphocytes and whole blood and estimated the PSI using rMATs^42^ using the same parameters described above. For each tissue, we identified samples with both STR genotypes and PSI to test for associations between STR copy number and PSI using the same linear regression model as used for the HBCC DLPFC analysis. We included the top 20 PEER factors, top 5 genotype PCs, sex, age and RNA-seq read counts as covariates. Notably, not all spliceSTRs detected in HBCC could be tested in GTEx. Of STRs detected as significant spliceSTRs in HBCC, 51.4% passed all quality filters in GTEx. The remainder failed primarily due to low call rate, likely driven by a high rate of PCR errors^42,100,101^ in the PCR+ GTEx WGS data (**Fig. S4c**). Of significant splicing events identified in HBCC, 70.53% were detected in at least one of the four GTEx tissues. This low overlap can be in part explained by tissue specificity^102^, as well as by the lower overall counts of splicing events detected in GTEx, driven by a combination of shorter read lengths^103^, lower read counts^104^ and lower sample sizes^42^ in GTEx (**Fig. S4**).

The other two studies compared^39,40^ used intron-based methods to identify splicing events, whereas we used the splice-event based splicing detection methods from rMATS^42^ (**Fig. S8a**). Additionally, while we imputed STRs using the SNP-STR reference haplotype panel^43^, the study by Hamanaka *et al*.^39^, which investigated splicing-TR associations, used GangSTR^88^ for STR genotyping and the study by Cui *et al.*^40^ on TR-xQTLs included a broad range of tandem repeats but excluded homopolymers (comprising ∼50% of STRs). These methodological differences further contributed to the limited overlap in the spliceSTRs tested across studies. To enable direct comparison, we identified matched intronic inclusion events from the other two studies. We downloaded the summary statistic data of 49 tissues from the splicing-TR (spl-TR) study^39^ (available for all tested STR-splice event pairs) and those of 8 brain tissues from the brain TR-xQTL study^40^ (available for tested pairs with FDR<5%). For each tissue, we compared the TR-splice events pairs to spliceSTRs identified in our study and selected those that: (1) shared the same STR, (2) exhibited comparable splicing events, defined by an intron length ratio (between the included intron in other studies and the intron between the two constitutive exons in our skipped-exon event) within the range of 0.95-1.05. We then tested the correlation of effect sizes across studies using Pearson’s *r*. Note, the same splicing event will show an opposite direction from the intron-based and rMATs^42^ based method. We flipped the direction of effect size from the other two studies using the intron-based method to make it easier to compare with those from our analysis.

### Fine-mapping spliceSTRs

We performed fine-mapping using three orthogonal methods. First, we performed model comparison using ANOVA to determine whether spliceSTRs for each splice event explained variation in splicing beyond a model consisting of the best SNP. For each splice event with a significant spliceSTR we defined the best SNP as that with the strongest *P*-value and then compared two linear models: Y* ∼ best_SNP (SNP-only model) vs. Y* ∼ best_SNP + spliceSTR (SNP + STR model) using the anova_lm function in the Python statsmodels.api.stats module. Y* represents the residues after regressing out all covariates (age, sex, RNA-seq read counts, PCs, PEER factors) from the PSI. We noticed many spliceSTRs with significant *P-*values from the ANOVA test show very small effect size on their associated splice events, those spliceSTRs may capture noises instead adding additional explanation beyond the best SNP to splicing variation. To be more stringent, we grouped all spliceSTRs by splice event and kept only STRs whose effect sizes are not significantly different (two-tailed Z-test, nominal *P*-values>0.05) from those of the STR with strongest *P* value at each event. After filtering, 107,670 out of 146,969 spliceSTRs remained. ANOVA *P*-values were adjusted using the fdrcorrection function in the statsmodels.stats.multitest module and the adjusted (under a FDR threshold of 10%) *P-*values were used for the determination of ANOVA-spliceSTRs (n = 17,524). A limitation of the ANOVA approach is that it only compares a single SNP and STR at a time. We therefore applied two additional methods: Lasso regression and Bayesian fine-mapping using FINEMAP.

Second, we performed Lasso regression to identify potential candidate variants that contribute to the variation of each significant splice event. For each significant splice event, PSI values were first adjusted by regressing out known covariates, including age, sex, RNA-seq read depth, genotype PCs, and PEER factors. We then selected all variants with nominal association *P*<0.05 from association testing as input predictors. On average, 145 variants per splice events passed this threshold and were included in the Lasso regression. Both the response (residual PSI) and predictor variables were standardized (*Z*-scored) prior to regression. To determine the optimal regularization parameter (α), we used five-fold cross-validation with the LassoCV function from the sklearn.linear_model module, specifying cv=5 and max_iter=5,000. A final Lasso model was then fitted using the Lasso function from the sklearn.linear_model module with the selected α and max_iter=5,000. The output coefficient for each variant from the Lasso model of each analyzed splice event was used for determining LS-spliceSTRs. We reasoned that non-causal variants should have coefficients close to 0 whereas variants that best explain the splicing event should have larger coefficients. However, estimated coefficients were rarely exactly 0 and instead followed a unimodal distribution centered around 0. Therefore, we grouped all tested variants by splice event and selected the spliceSTR whose STR had the largest absolute coefficient among all variants for each splice event as the Lasso-Selected spliceSTR (n = 7,830).

Third, we used FINEMAP^46^ v1.4.2 to prioritize spliceSTR signals out of all variants with nominal association *P*<0.05 for each splice event. On average, 145 variants per splice events passed this threshold and were included in the FINEMAP analysis. For each splice event, we constructed the input file required by FINEMAP including the effect size and the effect size’s standard error for all nominally associated variants. We set the values of maf, allele1 and allele2 columns to nan as that information is not required and many STRs tested have more than 2 alleles. To construct the LD input file for FINEMAP, pairwise LD between the STR and SNP was estimated using Pearson correlation between SNP dosages (0, 1, or 2) and STR dosages (sum of the best guess repeat lengths across both chromosome copies, in bp relative to the hg38 reference) across all samples. We ran FINEMAP v1.4.2 with non-default options --sss, --n-causal-snps 15, and --n-configs-top 100. The posterior inclusion probability (PIP) output for each variant for each splice event was used to determine FM -spliceSTRs. We selected spliceSTRs with posterior inclusion probabilities (PIPs) >=0.5 as FM-spliceSTRs (n = 4,647). Notably, for 64.60% (n = 15,866) of splicing events analyzed using FINEMAP, no single variant (SNP or STR) reached a threshold of PIP>0.5 (**Fig. S12**).

### Canonical repeat units and strand bias

For each STR, we defined the canonical repeat unit as the lexicographically first repeat unit when considering all rotations and strand orientations (relative to coding strand) of the repeat sequence. For example, the canonical repeat unit for the repeat sequence CAGCAGCAGCAG would be AGC. We further defined the strand bias of repeat units based on their nucleotide composition and repeat unit length. 9,628 repeat units with equal numbers of “A” and “T” nucleotides and 3,234 repeat units in which “G” and “C” bases account for at least 50% of the sequence were excluded in the strand bias analysis. Finally, a total of 242,423 repeat units in which “A” and “T” bases together account for at least 50% of the sequence were used for strand bias analysis. These repeats were further grouped according to their relative “T” and “A” content, with repeat units containing more “T” classified as “T” rich repeat units.

### Enrichment analyses

Enrichment analyses were performed using a two-sided Fisher’s exact test as implemented in the fisher_exact function of the Python package scipy.stats v1.5.2. The overlap of STRs with each annotation was performed using the intersectBed tool of the BEDTools^105^ suite v2.30.0. Genomic annotations were obtained by downloading custom tables using the UCSC Genome Browser^94^ table browser tool to select genes and gene predictions using the GENCODE V38^106^ track. Transcription start (TSS) and end sites (TES) were determined by grouping all transcripts by gene and selecting the combination that spanned the longest range between TSS and TES for each gene. The distances of STRs to the transcription start site (TSS) and transcription end site (TES) were stranded based on the gene’s transcriptional direction. These distances were then scaled relative to the total length between the associated gene’s TSS and TES. Scaled values less than 0 indicate positions upstream of the TSS, values greater than 1 indicate positions downstream of the TES, and values between 0 and 1 represent positions within the transcribed region.

Gene-set over-representation analyses were performed separately using PANTHER v19.0^107–109^ and ShinyGO v0.85.1^110–113^ for three gene sets: genes associated with all spliceSTRs, genes associated with leniently fine-mapped spliceSTRs, and genes associated with strictly fine-mapped spliceSTRs. For each analysis, all genes tested in the spliceSTR association analysis were used as the background gene set. In PANTHER, over - representation analysis was performed using the PANTHER Pathways annotation set (a curated collection of signaling and metabolic pathways) and Fisher’s exact test, with correction for multiple testing using the false discovery rate (FDR). In ShinyGO, Kyoto Encyclopedia of Genes and Genomes (KEGG) pathway enrichment was assessed using the hypergeometric test, with *P*-values adjusted using the Benjamini–Hochberg FDR procedure. Pathways with FDR<0.05 were considered significantly enriched.

### RBP enrichment and binding association analysis

Transcriptome-wide RNA-protein interactions peaks data were downloaded from the study by Boyle *et al*.^48^, which reanalyzed the crosslinking and immunoprecipitation followed by sequencing (CLIP-seq) dataset^58^ using the Skipper^48^ pipeline. Reproducible enriched binding windows for the K562 and HepG2 cell lines were downloaded and intersected with all STRs tested in our analysis, requiring strand matching. For each RBP, the STR was labeled as overlapping if it overlapped a peak detected in either cell line. We also downloaded the bigWig files generated from the CLIP-seq datasets^58^ to visualize raw RBP binding signal profiles at nucleotide resolution around spliceSTR loci.

To evaluate whether variation in STR length influences RBP binding, we first identified STRs that overlapped RBP binding peaks, again requiring strand concordance. To control for biases due to STR length affecting overlap detection, all STRs were centered and uniformly extended to a fixed window of 100 base pairs. STRs were then grouped by repeat unit sequence. For each repeat unit-RBP pair, we tested for associations between repeat unit copy number and the presence of overlap with RBP binding sites using logistic regression. Specifically, we modeled the probability of overlap using the following logistic regression equation: 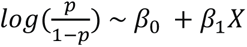 where *p* is the probability that an STR overlaps an RBP binding site and *X* is the length of the STR. The binary response variable was defined as 1 if the repeat unit overlapped a binding site and 0 otherwise. We required each RBP-unit pair to have at least 100 overlapping loci, and at least 10 unique STRs in each length of the tested repeat unit. Logistic regression was performed using the logit function from the Python statsmodels.api module, which estimates regression coefficients via maximum likelihood and reports two-tailed *P*-values based on Wald *z*-statistics.

### Colocalization of fine-mapped spliceSTRs with published GWAS signals

Schizophrenia GWAS signals were downloaded from the study by Trubetskoy *et al.*^70^. Autism spectrum disorders GWAS signals were downloaded from the study by Grove *et al.*^114^. Bipolar disorder GWAS signals were downloaded from the study by Mullins *et al.*^115^. Attention deficit hyperactivity disorder GWAS signals were downloaded from the study by Demontis *et al.*^116^. Major depressive disorder GWAS signals were downloaded from the study by Howard *et al.*^117^. The GWAS summary statistics for schizophrenia (PGC3_SCZ_wave3.primary.autosome.public.v3.tsv.gz), autism spectrum disorders (iPSYCH-PGC_ASD_Nov2017.gz), bipolar disorder (daner_bip_pgc3_nm_noukbiobank.txt), attention deficit hyperactivity disorder (ADHD_meta_Jan2022_iPSYCH1_iPSYCH2_deCODE_PGC.meta), and major depressive disorder (MDD_jamapsy_Giannakopoulou_2021_exclude_whi_23andMe.txt) were downloaded from the Psychiatric Genomics Consortium (PGC) website. Alzheimer’s disease GWAS signals were downloaded from Bellenguez *et al.*^118^ and the summary statistics were downloaded from the GWAS catalog (study ID: GCST90027158). For datasets mapped to hg19 coordinates, we lifted the coordinates to hg38 using LiftOver^93^ with a chain file downloaded from the UCSC Genome Browser^94^.

LD between the STR and their nearby lead GWAS SNP was estimated using all European samples from 1000 Genomes Project by calculating the Pearson correlation between SNP dosages (0, 1, or 2) and STR dosages (hard-called total repeat lengths across both chromosome copies, measured in base pairs relative to the hg38 reference genome) genotyped from our previous study^44^.

Colocalization analysis between fine-mapped spliceSTRs and GWAS signals was performed using the coloc.abf function from the coloc^119^ R package v5.2.3. For all traits, two datasets were provided as input: Dataset 1 included SNP effect sizes and their variances from the splicing association tests. Since PSI values for splicing events were Z-scaled, this dataset was specified with type = “quant” and sdY = 1. Dataset 2 included effect sizes and their variances from GWAS summary statistics for each trait, with type = “CC” (case-control), along with the proportion of cases for each trait.

### Identifying spliceSTRs overlapping STRs previously associated with brain-related disorders

Candidate repeat expansion loci for ASD were obtained from Trost *et al.*^73^, and those for schizophrenia were obtained from Mojarad *et al.*^72^. Because these loci were identified using ExpansionHunter Denovo^120^, which reports approximate repeat-expansion coordinates rather than precise STR boundaries, we defined potentially matching loci using two criteria: (1) the reported locus and the STR in our spliceSTR catalog shared the same repeat motif, and (2) their start coordinates differed by less than 100 bp. For each potentially matching locus, we determined whether the STR was significantly associated with a splicing event in our dataset (FDR < 5%) and whether the spliceSTR was supported by our lenient or strict fine-mapping analyses.

For Alzheimer’s disease, we compared our spliceSTR catalog with STR associations reported in a recent genome-wide association study from Gmelin *et al.*^75^. Because that study represented STR genotypes in a bi-allelic format and used a different STR reference panel, the reported STR coordinates could differ slightly from those in our catalog. We therefore defined potentially matching loci as STRs that shared the same repeat motif and whose start coordinates matched after rounding down to the nearest 10 bp. A complete list of matched Alzheimer’s disease-associated STRs reaching genome-wide significance (*P*<5 × 10⁻⁸) and their corresponding splicing associations is provided in **Supplementary Table 2**.

## Declarations

### Ethics approval and consent to participate

Not applicable.

### Consent for publication

All data used in this study were obtained through dbGaP (phs000979.v3.p2) under an approved data access request. Participants in the original studies have been consented for general research use, including public posting of genomic summary results.

### Availability of data and materials

STR genotype calls for GTEx samples are available on Anvil at accession GTEx_v8_Hipstr_calls. HBCC and GTEx spliceSTR summary statistics are provided in **Supplementary Datasets 1-2** available on Figshare: https://figshare.com/s/f38632f668e232d9c9bb. Custom code for generating figures is available at https://doi.org/10.5281/zenodo.21712391.

Additional datasets and resources used in this study are listed below:

SNP-STR reference haplotype panel: https://gymreklab.com/2018/03/05/snpstr_imputation.html

Beagle genetic maps: https://bochet.gcc.biostat.washington.edu/beagle/genetic_maps/plink.GRCh37.map.zip

Liftover chain file: https://hgdownload.cse.ucsc.edu/goldenpath/hg19/liftOver/hg19ToHg38.over.chain.gz

ENCODE RNA-seq pipeline: https://github.com/ENCODE-DCC/long-rna-seq-pipeline/raw/master/DAC/STAR_RSEM.sh

GRCh38.p13 reference genome: https://ftp.ebi.ac.uk/pub/databases/gencode/Gencode_human/release_38/GRCh38.p13.genome.fa.gz

GENCODE v38: https://ftp.ebi.ac.uk/pub/databases/gencode/Gencode_human/release_38/gencode.v38.annotation.gtf.gz

Statmodels package: https://www.statsmodels.org

HipSTR hg38 reference panel: https://github.com/HipSTR-Tool/HipSTR-references/raw/master/human/hg38.hipstr_reference.bed.gz

Scipy.stats package: https://docs.scipy.org/doc/scipy/reference/stats.html

Fusera: https://github.com/ncbi/fusera

RBP peaks: https://figshare.com/articles/dataset/Skipper_RNA-protein_interaction_profiles/21206009

Psychiatric Genomics Consortium summary statistic downloads: https://pgc.unc.edu/for-researchers/download-results/

RBP bigWig files: https://www.encodeproject.org/search/?type=Experiment&control_type!=*&status=released&perturbed=false&assay_title=eCLIP

### Authors’ contributions

M.G. and A.G. conceived the study, supervised analyses, and wrote the manuscript. Y.L. led, designed and performed the analyses and wrote the manuscript. J.M. provided guidance on the FINEMAP analysis.

## Acknowledgements

We acknowledge Yuren Dong, Ziyang Zhang, and Daniela Nachmanson who helped conceive the study and develop preliminary analysis pipelines. We thank Tara Mirmira for proofreading the manuscript and providing helpful comments. We additionally acknowledge Gene Yeo and Charlene (Hsuan-lin) Her for advice on analysis of RBP datasets.

HBCC data was accessed through the Database of Genotypes and Phenotypes (dbGaP) accession phs000979.v3.p2 which was supported by the Intramural Research Program of the NIMH(NCT00001260, 900142).

The Genotype-Tissue Expression (GTEx) Project was supported by the Common Fund of the Office of the Director of the National Institutes of Health (commonfund.nih.gov/GTEx). Additional funds were provided by the NCI, NHGRI, NHLBI, NIDA, NIMH, and NINDS. Donors were enrolled at Biospecimen Source Sites funded by NCI\Leidos Biomedical Research, Inc. subcontracts to the National Disease Research Interchange (10XS170), Roswell Park Cancer Institute (10XS171), and Science Care, Inc. (X10S172). The Laboratory, Data Analysis, and Coordinating Center (LDACC) was funded through a contract (HHSN268201000029C) to The Broad Institute, Inc. Biorepository operations were funded through a Leidos Biomedical Research, Inc. subcontract to Van Andel Research Institute (10ST1035). Additional data repository and project management were provided by Leidos Biomedical Research, Inc.(HHSN261200800001E). The Brain Bank was supported supplements to University of Miami grant DA006227. Statistical Methods development grants were made to the University of Geneva (MH090941 & MH101814), the University of Chicago (MH090951,MH090937, MH101825, & MH101820), the University of North Carolina - Chapel Hill (MH090936), North Carolina State University (MH101819), Harvard University (MH090948), Stanford University (MH101782), Washington University (MH101810), and to the University of Pennsylvania (MH101822). The datasets used for the analyses described in this manuscript were obtained from dbGaP at http://www.ncbi.nlm.nih.gov/gap through dbGaP accession number phs000424.v8.p2.

