## Supplemental Figures for "The contribution of short tandem repeats to splicing variation in the human prefrontal cortex"

### Supplementary Figures

#### Supplementary Figure 1

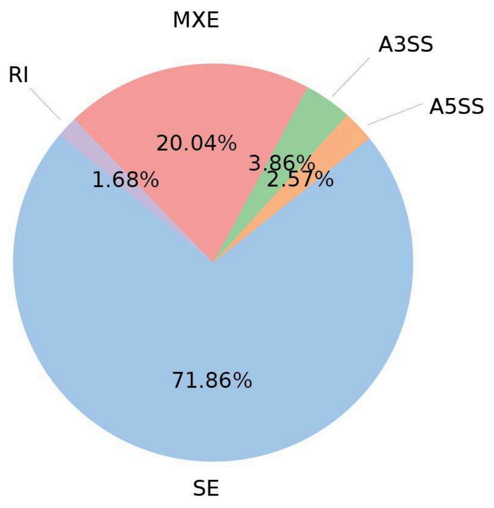


**Summary of splice event types identified by rMATs in the HBCC dataset.** A total of 548,614 splice events were identified by rMATs. The pie chart shows the percentage of the five alternative splicing patterns considered: skipped exon (SE, n=394,222), alternative 5' splice sites (A5SS, n=14,094), alternative 3' splice sites (A3SS, n=21,171), mutually exclusive exons (MXE, n=109,921), or retained intron (RI, n=9,206).

### Supplementary Figure 2

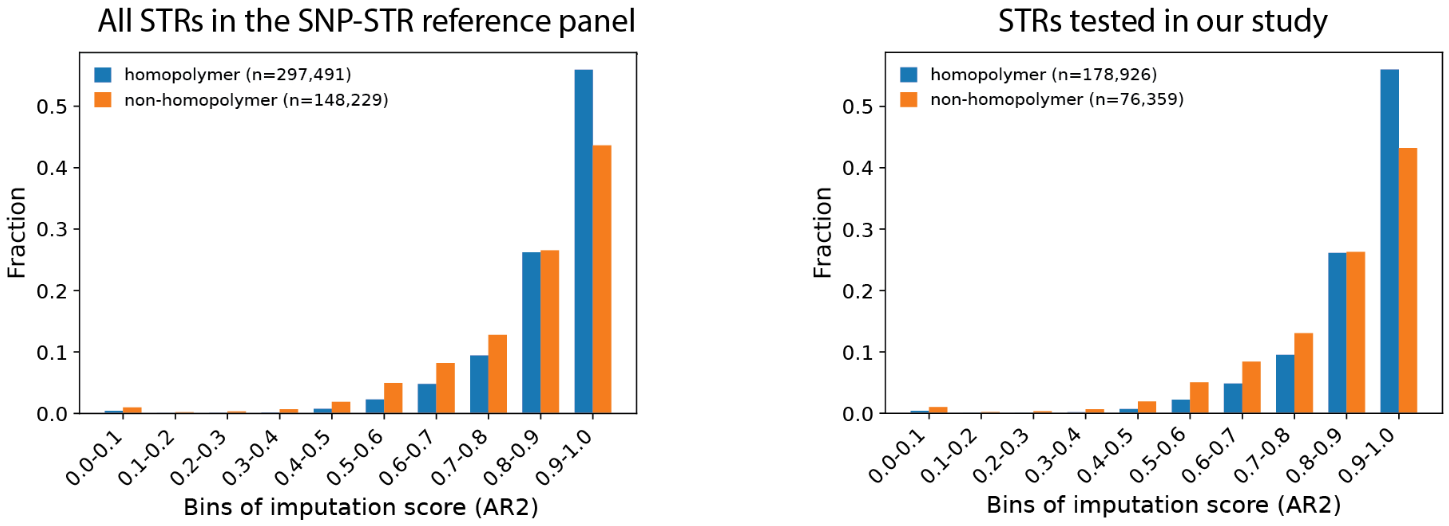


**Distribution of imputation quality scores for homopolymer and non-homopolymer STRs.** STRs were grouped by imputation quality scores bins (x-axis), and bar heights represent the proportion of loci in each bin within each repeat class (y-axis). The total number of homopolymer (blue) and non-homopolymer (orange) STRs is shown in the legend. Imputation scores were obtained from the AR2 field of the SNP-STR reference haplotype panel. The left panel shows all STRs from the SNP-STR reference panel, and the right panel shows STRs tested in our spliceSTR association analysis.

Supplementary Figure 3

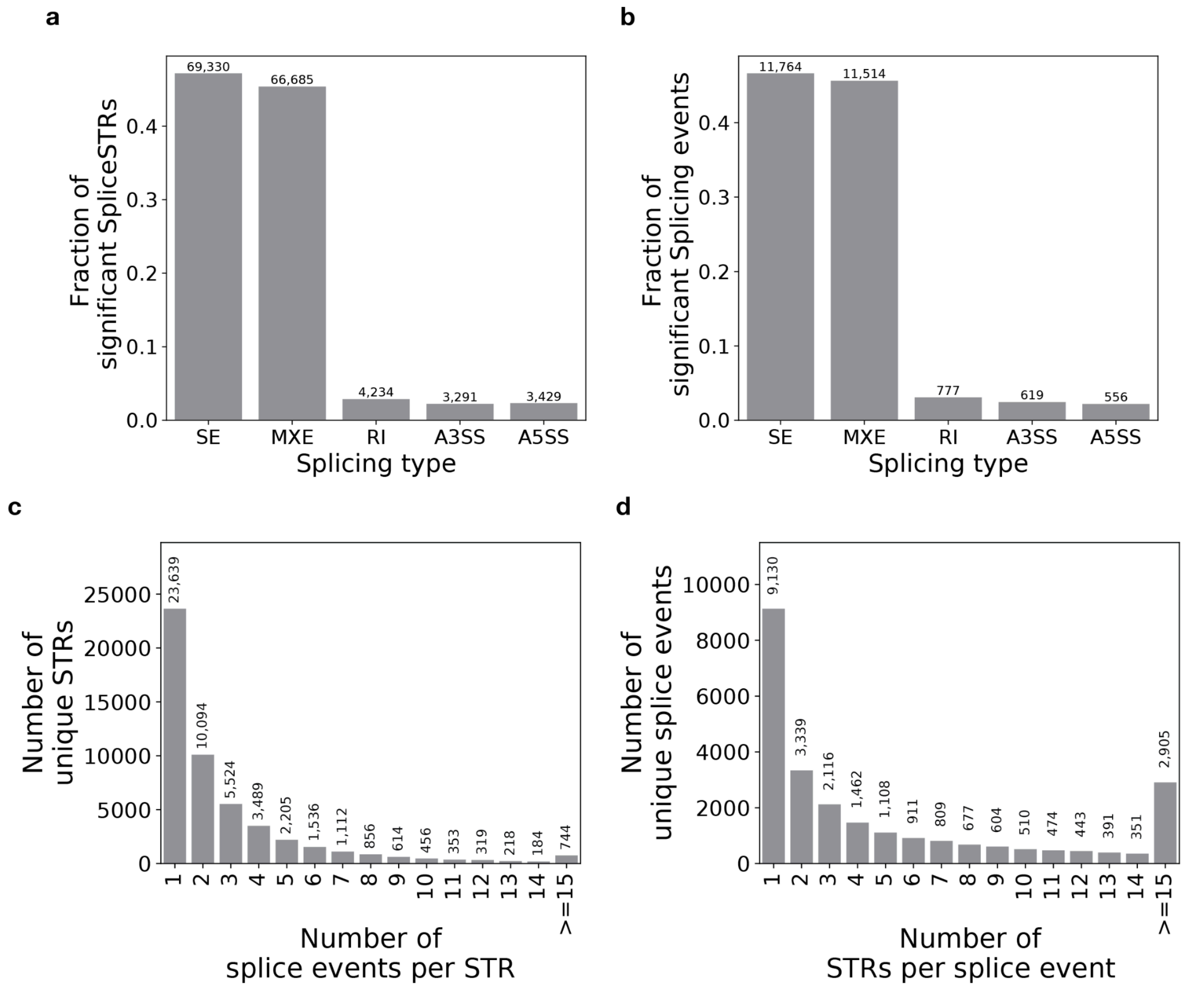


**Summary of splicing event types across significant spliceSTRs. (a) The fraction of STR-splice event pairs tested that were identified as significant spliceSTRs (FDR 5%) stratified by splice event type.** A total of 146,969 significant STR-splice event pairs were identified. **(b) The fraction of tested splicing events identified as having at least one significant spliceSTR stratified by event type.** A total of 25,230 unique splicing events had at least one spliceSTR. In each panel, the *x*-axis represents the type of splice event. Exact values are shown above each bar. **(c) The number of unique splice events significantly associated per STR. A total of 51,343 unique significant STRs were identified. (d) The number of unique STRs significantly associated per splice event.** A total of 25,230 unique significant splice events were identified. Bars represent counts of unique STRs or splice events (*y*-axis) across different categories (*x*-axis). Numerical values above each bar indicate the total count for that category. The median number of unique splice events per STR is 2 and the median number of unique STRs per splice event is 3.

### Supplementary Figure 4

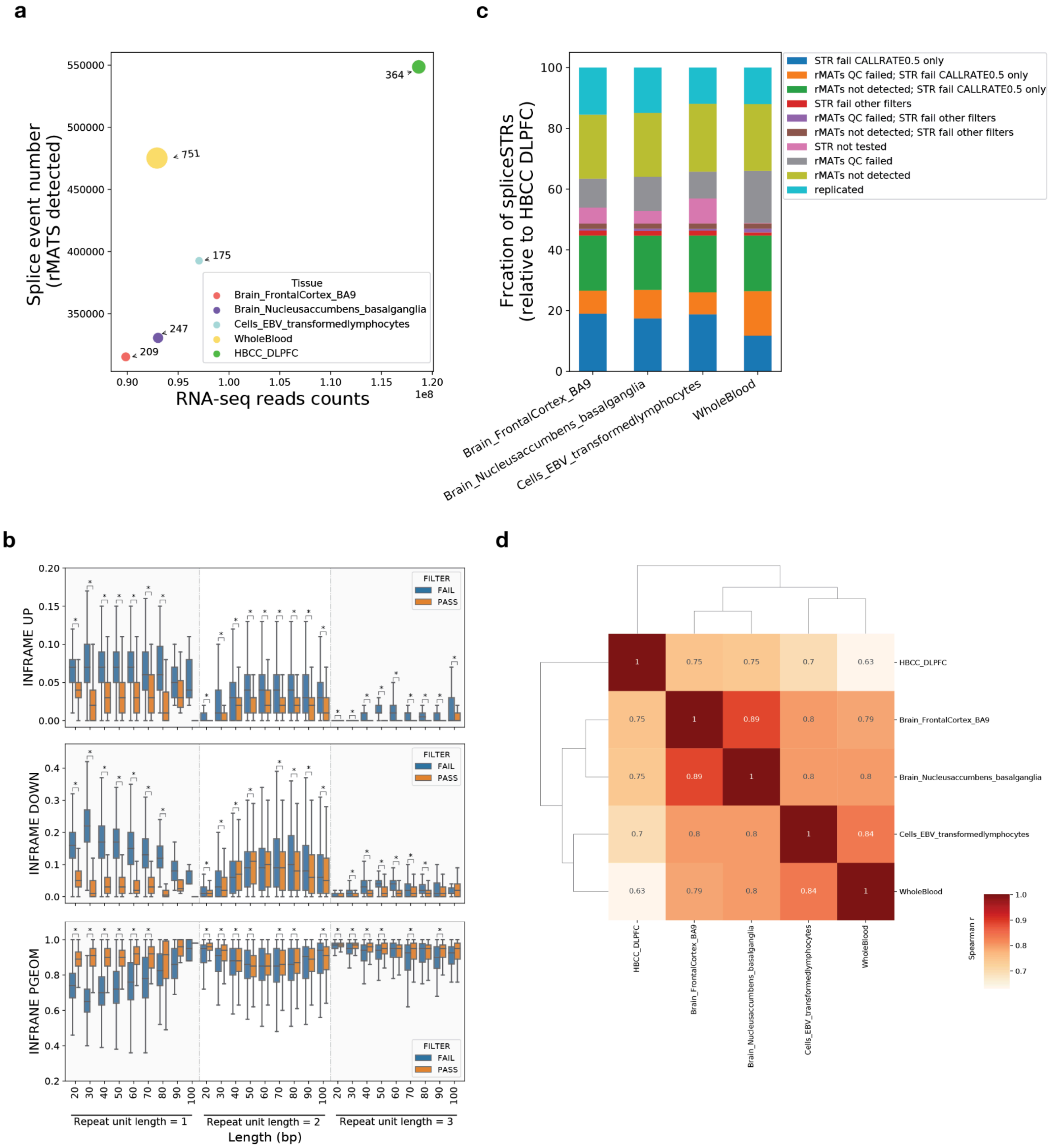


**Comparison of spliceSTRs identified in the HBCC DLPFC with those found in four GTEx tissues. (a) Relationship between read counts from RNA-sequencing and number of splice events detected by rMATs.** The *x*-axis shows the mean read counts of the RNA-sequencing across all samples in each tissue. The *y*-axis shows the total number of alternative splice events detected by rMATs across all samples per tissue. Each dot represents a single tissue as indicated by the legend. The size of the dot corresponds to the sample size of each tissue. The numbers show the exact sample size of each tissue. Although HBCC DLPFC does not have the largest sample size, it has the deepest RNA-seq coverage and correspondingly the highest number of splice events detected by rMATs. **(b) Comparison of stutter model parameters between STRs that passed and failed HipSTR locus-level filters** (**Methods**). Per-locus stutter parameters were estimated from PCR+ high-coverage WGS for GTEx samples using HipSTR. Box plots in each row show the distribution of the three stutter parameters output by HipSTR for each STR (top=INFRAME_UP, which denotes the estimated fraction of reads showing a PCR stutter error that increases repeat length; middle=INFRAME_DOWN, which denotes the estimated fraction of reads showing a PCR stutter error that decreases repeat length; and bottom=INFRAME_PGEOM, which denotes the fraction of stutter errors that result in changes of a single repeat unit). Results are stratified by repeat unit length. Horizontal lines show median values, boxes span from the 25th percentile (Q1) to the 75th percentile (Q3). Whiskers extend to $Q1 - 1.5 \times IQR$ (bottom) and $Q3 + 1.5 \times IQR$ (top), where IQR is the interquartile range (Q3–Q1). The *x*-axis in each panel shows bins of STR total length in base pairs. The color indicates "PASS" or "FAIL" by HipSTR filters (see **Methods**). The overall distributions of stutter parameters are similar to those observed in PCR+ high-coverage WGS samples from the H3Africa cohort[1]. Asterisks denote statistical significance (Mann–Whitney U test, nominal $P < 0.05$) between STRs that passed vs. failed HipSTR call filters. **(c) Categorization of spliceSTR events identified as significant in HBCC DLPFC that could be tested in each of the four GTEx tissues considered.** The *x*-axis shows the name of the tissue from GTEx. The *y*-axis shows the percentage of significant spliceSTRs detected in HBCC DLPFC in each category. Cyan indicates spliceSTRs for which both the STR and splicing event passed QC filters in the GTEx dataset. Other colors indicate the percentage of spliceSTRs filtered due to failing either STR genotyping or rMATs QC steps. **(d) Pairwise correlation of effects of significant spliceSTRs across tissues.** Each cell shows the Pearson correlation between the effect sizes of spliceSTRs identified in either of the paired tissues at FDR 5%.

### Supplementary Figure 5

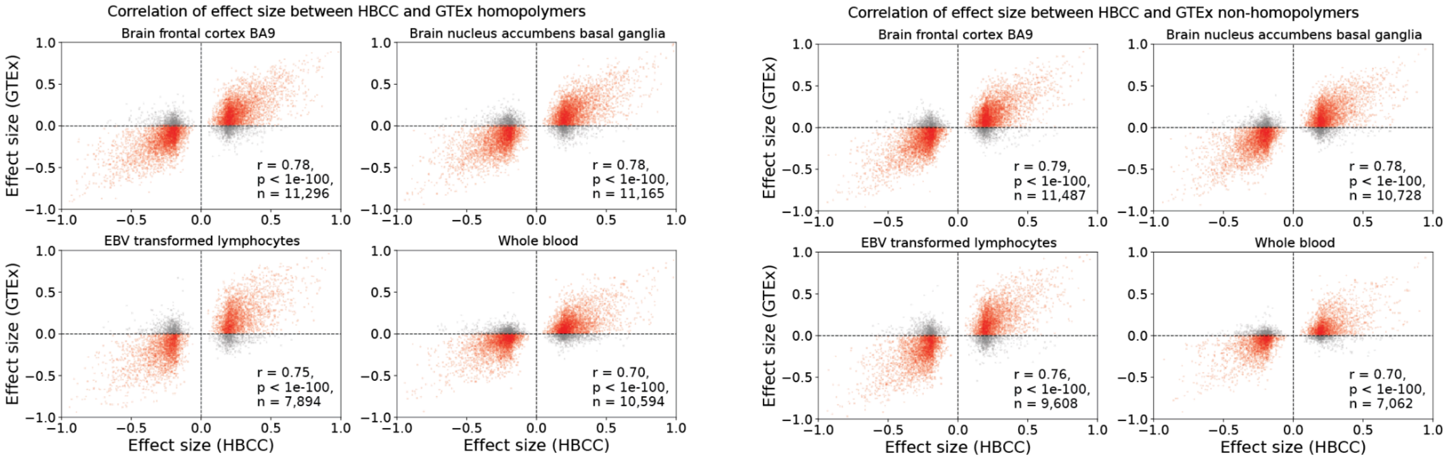


**Comparison of spliceSTR effect sizes stratified by repeat unit length across cohorts.** The *x*-axis of each plot represents effect sizes measured in HBCC. The *y*-axis of each plot represents effect sizes measured in GTEx. The title of each subplot denotes the tissue from GTEx. Only spliceSTRs reaching FDR<5% in HBCC are included. The Pearson's correlation (*r*), two-tailed *P*-value, and number of spliceSTRs compared (*n*) are annotated in each panel. The left set of panels shows results for homopolymer spliceSTRs and the right set of panels shows results for non-homopolymer spliceSTRs.

### Supplementary Figure 6

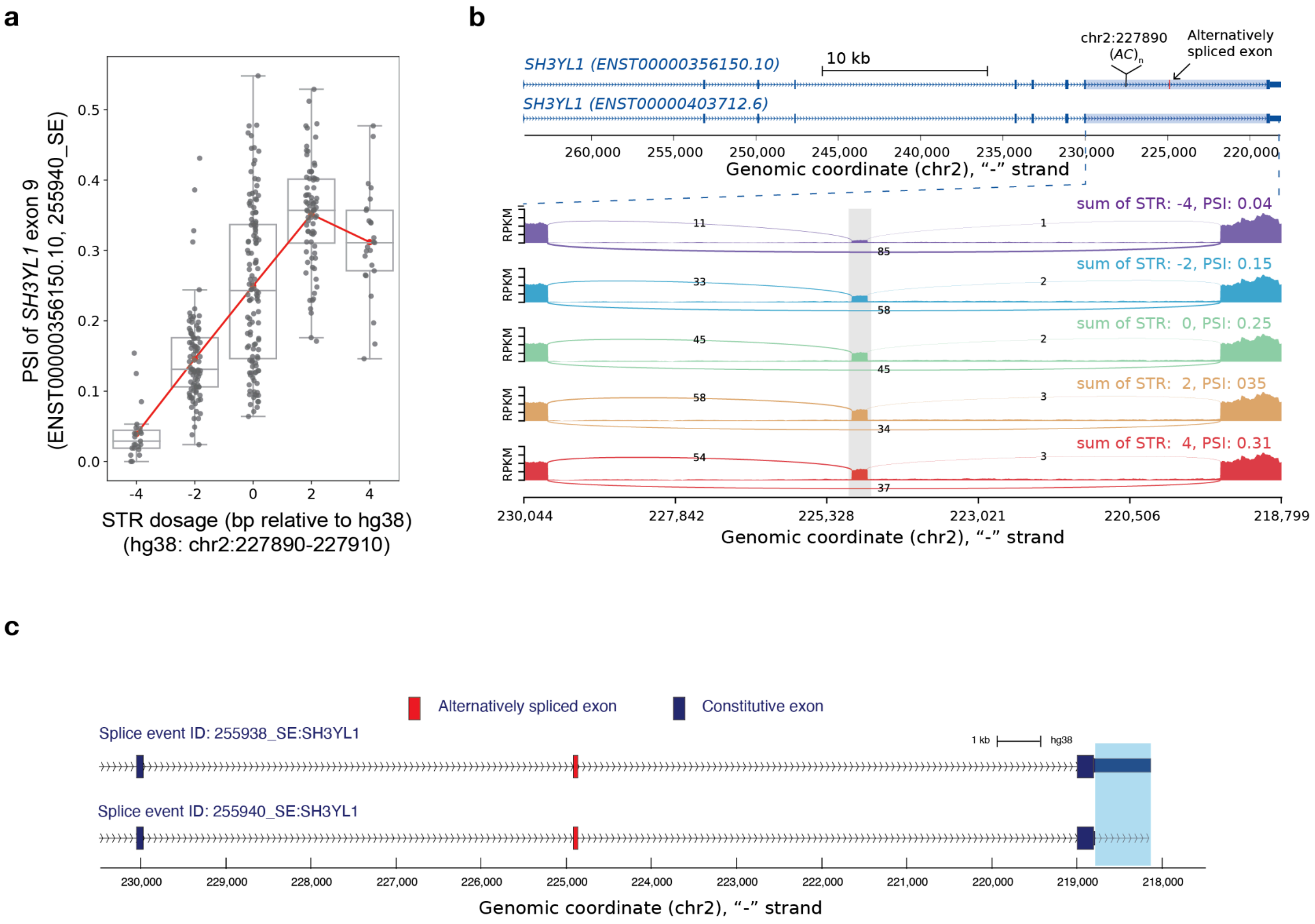


**Example spliceSTR identified in the HBCC DLPFC that is also significant in three of four GTEx tissues tested. (a) Example spliceSTR.** The *x*-axis represents the STR dosage (sum of repeat lengths across both chromosome copies, in bp relative to the hg38 reference) of an STR (hg38:chr2:227890-227910) within an intron of *SH3YL1*. The *y*-axis represents the PSI of exon 9 (ENST00000356150.10) and the splice event ID (225940_SE). Each dot represents a single individual. Box plots summarize the distribution of PSI values. Horizontal lines show median values, boxes span from the 25th percentile (Q1) to the 75th percentile (Q3). Whiskers extend to Q1 − 1.5 × IQR (bottom) and Q3 + 1.5 × IQR (top), where IQR is the interquartile range (Q3–Q1). The red line shows the mean expression for each x-axis value. **(b) Sashimi plot showing read-level evidence for the example spliceSTR.** This spliceSTR has the same STR and associated exon as that shown in **Fig. 1d**, but was identified in GTEx with a different boundary of the downstream constitutive exon (see panel **c**). The full exonic structure of two representative *SH3YL1* transcripts and the location of the splicesSTR (hg38:chr2:227890-227910) are shown in the top panel. Samples were grouped by STR dosage. The alternatively spliced exon is marked by a grey box. To better visualize the spliced exon, intronic regions are displayed at one-fifth the scale of exonic regions. Coordinates are in descending order since annotations are shown for the reverse strand relative to hg38. Sashimi plots were generated using rmats2sashimiplot (v2.0.4). **(c) Exonic structure of the two similar splice events.** The exonic structure of splice event ID 255938_SE:SH3YL1 identified in HBCC from **Fig 1d** is shown in the top panel. A similar splice event with ID 255940_SE:SH3YL1 identified in both HBCC and GTEx is shown in the bottom panel. The boundary difference between the two splice events is highlighted with a blue box.

### Supplementary Figure 7

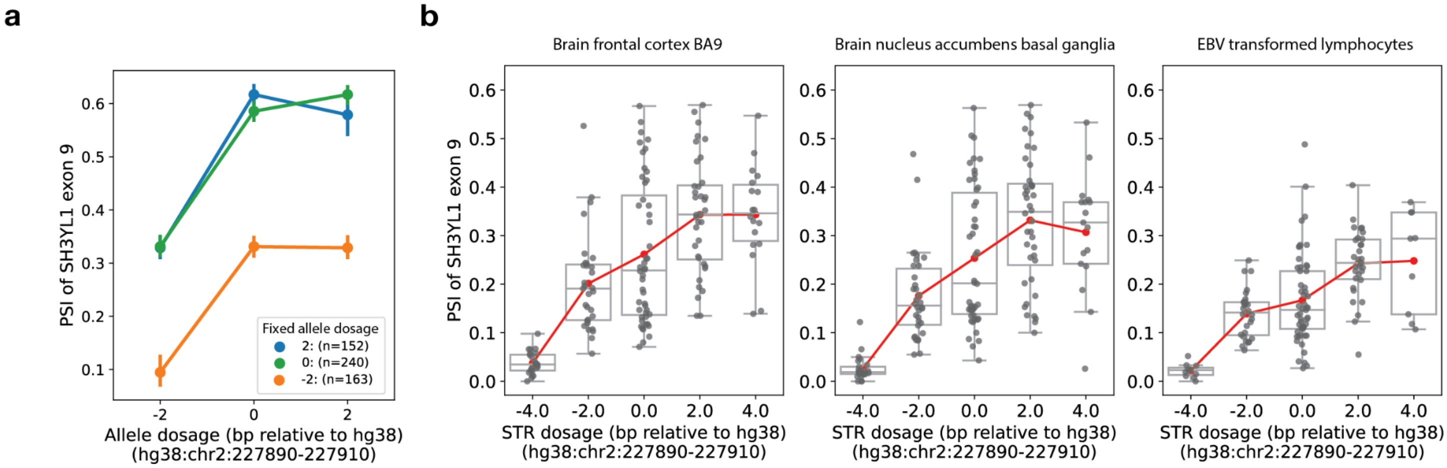


**Evaluation of non-linearity at the *SH3YL1* spliceSTR example. (a) *SH3YL1* exon 9 splicing association stratified by associated STR allele length.** The *x*-axis indicates the dosage (length) of one STR allele, in base pairs relative to the hg38 reference, while the other allele is held constant. The STR is located at hg38 chr2:227890–227910 within an intron of *SH3YL1*. The *y*-axis represents the PSI of exon 9 (ENST00000356150.10). Each line represents a group that was stratified by the dosage of the fixed allele and indicated by different colors (orange, −2 bp; green, 0 bp; blue, +2 bp relative to the hg38 reference). The points indicate mean PSI and error bars indicate confidence intervals of 95%. **(b) Replication of the *SH3YL1* exon 9 splicing association across GTEx tissues.** We focused on three tested GTEx tissues (Brain frontal cortex, Brain nucleus accumbens basal ganglia and EBV transformed lymphocytes), for which the same STR-splice event pair was also identified as significant significant (FDR<5%); The *x*-axis represents the STR dosage (sum of repeat lengths across both chromosome copies, in bp relative to the hg38 reference) of the STR located at chr2:227890-227910 (hg38), which is within an intron of *SH3YL1*. The *y*-axis represents the PSI of exon 9 (ENST00000356150.10). Each dot represents a single individual. Box plots summarize the distribution of PSI values. Horizontal lines show median values, boxes span from the 25th percentile (Q1) to the 75th percentile (Q3). Whiskers extend to Q1 − 1.5 × IQR (bottom) and Q3 + 1.5 × IQR (top), where IQR is the interquartile range (Q3–Q1). The red line shows the mean expression for each x-axis value. Tissue names are indicated above each panel.

### Supplementary Figure 8

**a**

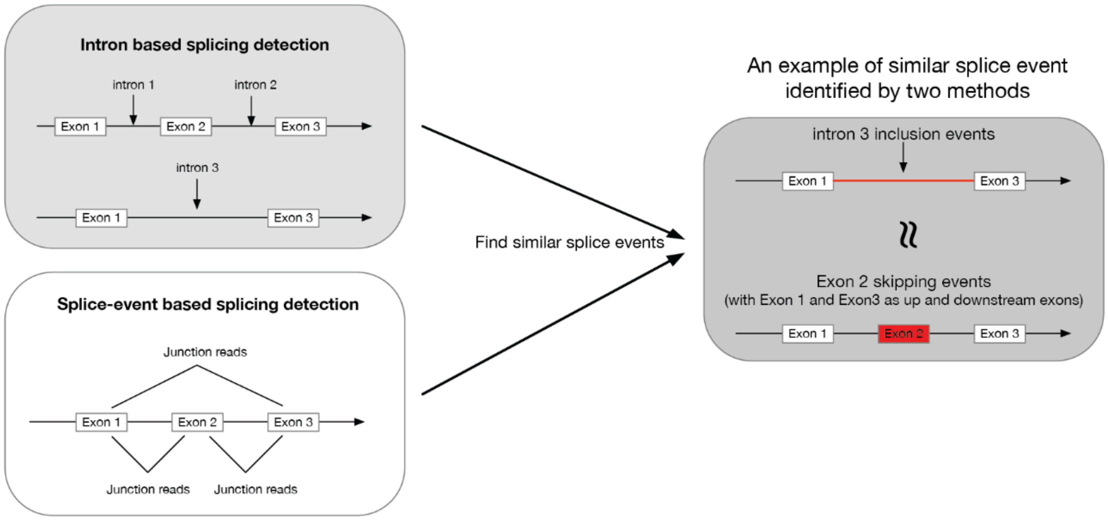


**b**

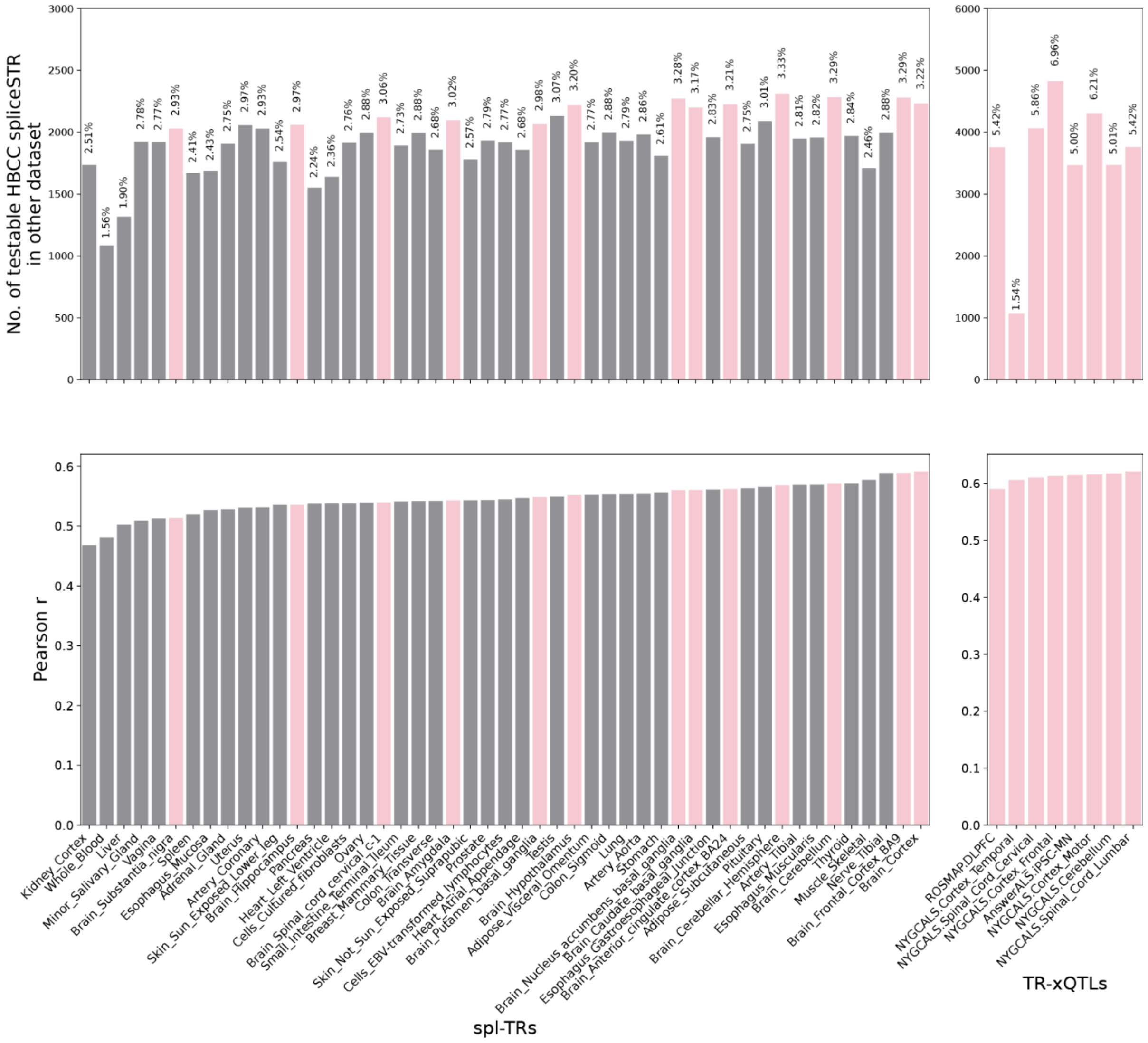

**Comparison of spliceSTRs identified in HBCC DLPFC with those reported in other studies. (a) Schematic diagram shows the two distinct methods for detecting splice events.** Intron-based detection measures the inclusion of an intron, whereas splice-event based detection assesses the inclusion of an exon. Our study used rMATs to perform splice-event based detection, whereas the studies compared to used intron-based detection. To enable comparison, we focused on skipped exon events identified by rMATS and matched them to corresponding intron-based events by comparing intron lengths. Notably, exon inclusion corresponds to the exclusion of the intron between the two flanking constitutive exons. **(b) Replication of HBCC DLPFC spliceSTRs vs. other studies (spl-TRs study[2], TR-xQTLs study[3]).** The upper panel shows the number of significant skipped exon spliceSTRs identified in HBCC DLPFC (FDR<5%) for which summary statistics were available from other studies. Hamanaka et al. (spl-TRs) provided summary statistics for all tested STR-splice event pairs in their study, whereas the TR-xQTL study provided summary statistics only for significant spliceSTRs (FDR<5%). The percentage of events that could be tested in the other datasets is labeled above each bar. The lower panel displays the Pearson correlation of effect sizes for shared skipped exon spliceSTRs between HBCC DLPFC and each tissue from the other studies. Tissue names from each study are shown on the *x*-axis. Pink and grey bars represent brain and non-brain tissues respectively.

### Supplementary Figure 9

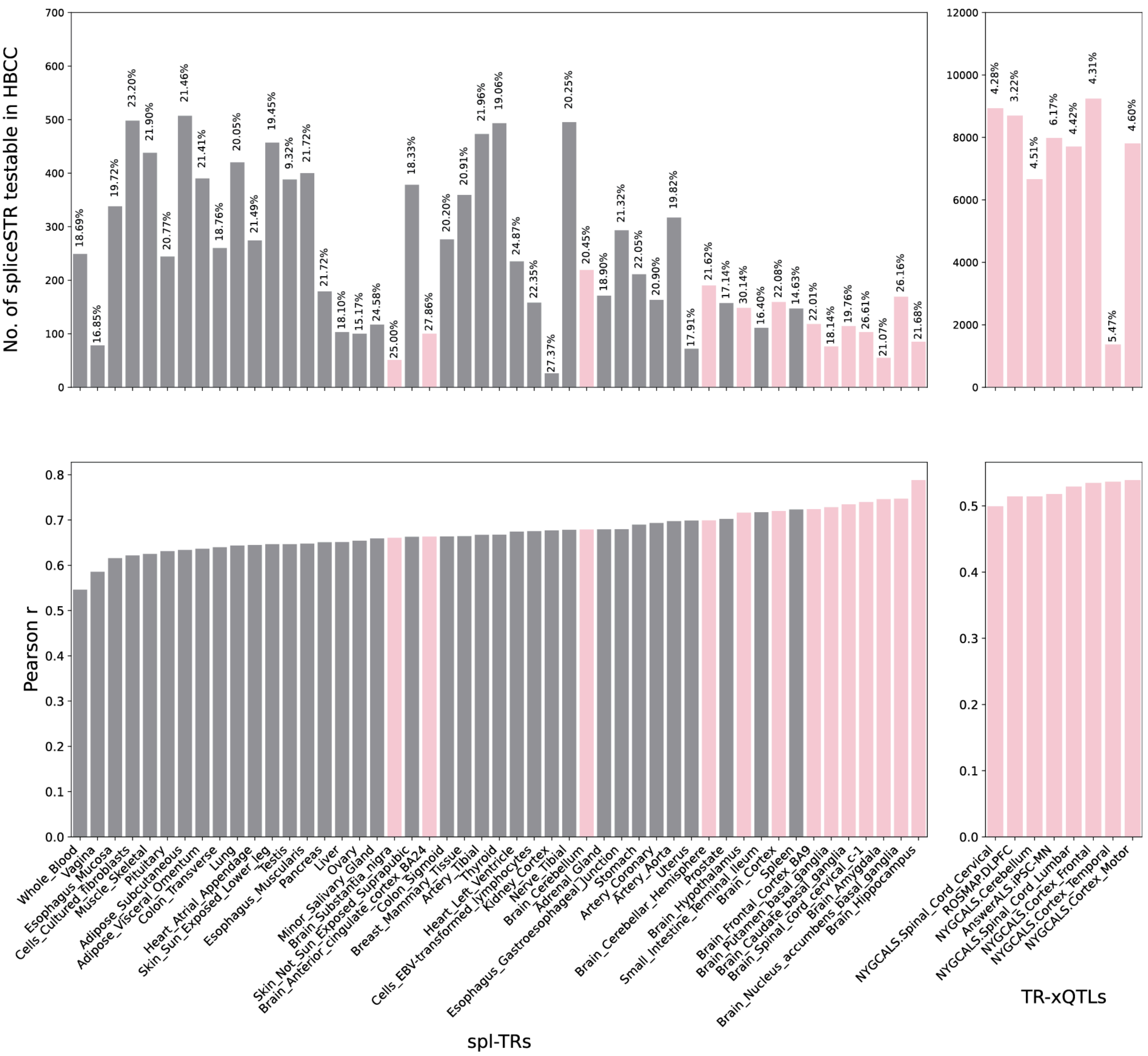


**Replication of spliceSTRs identified by other studies in the HBCC DLPFC dataset.** The upper panel shows the number of spliceSTRs identified by other studies (in both cases at FDR<5%) that were tested in our HBCC dataset. Bars are labeled with the percentage of total significant events that were tested in our study. The lower panel displays the Pearson correlation of effect sizes across these overlapping splice event-STR pairs. Tissue names from each study are shown on the *x*-axis. Pink and grey bars represent brain and non-brain tissues respectively.

### Supplementary Figure 10

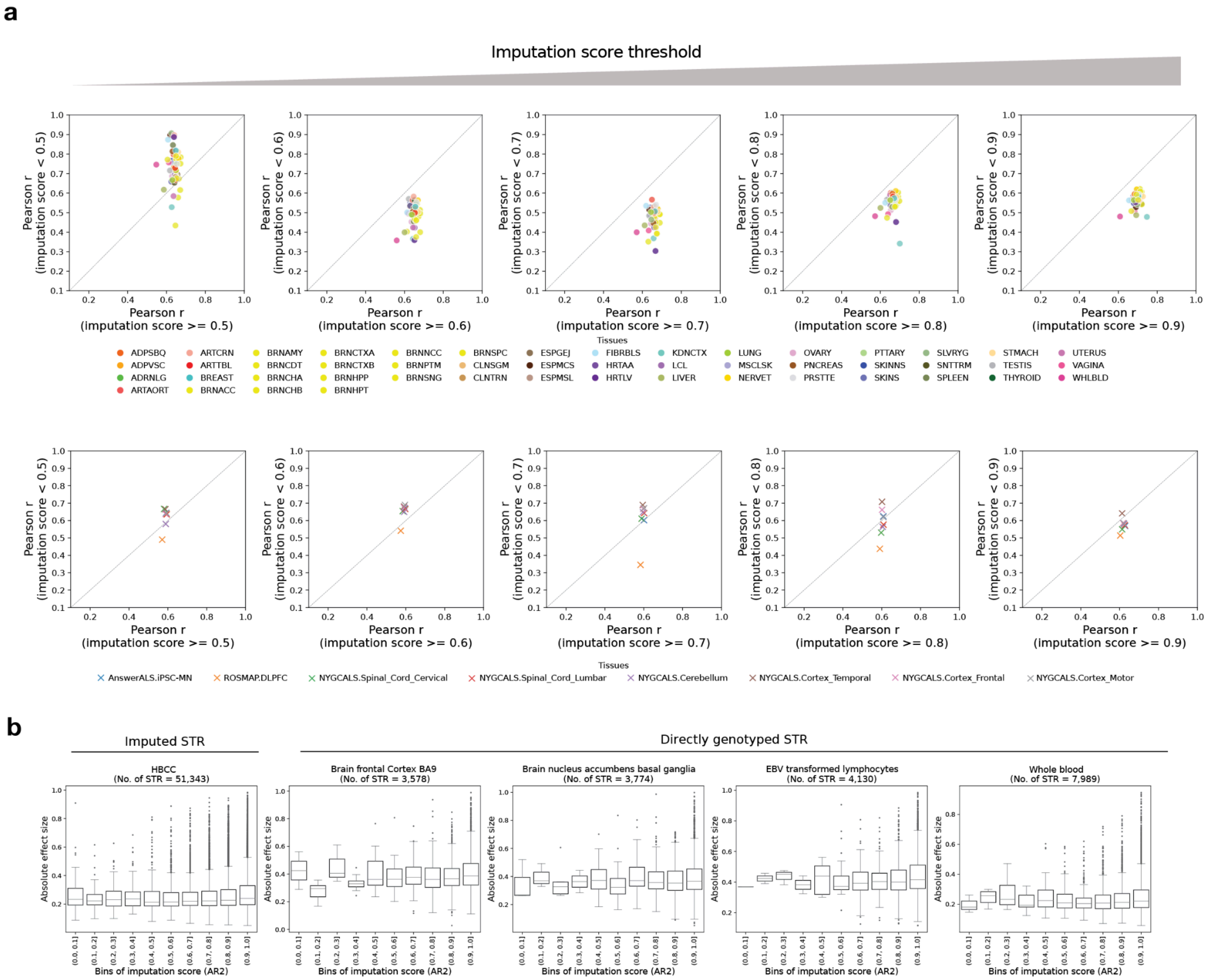


**Evaluating the relationship between STR imputation quality and spliceSTR replication and effect sizes. (a) Correlation of effect sizes across this and other studies stratified by imputation score.** Top row = HBCC (this study) vs. effect sizes inferred using GTEx data by Hamanaka et al. Each dot represents a single tissue. Plots show the Pearson correlation (r) of spliceSTR effect sizes when considering STRs with scores at or above (*x*-axis) vs. below (*y*-axis) a particular STR imputation score (annotated on the *x*-axis label). Bottom row=HBCC (this study) vs. effect sizes inferred from brain datasets analyzed in Cui et al. Each dot represents a different brain dataset. Plot elements are similar to the top row. Gray lines give the x=y diagonal. Imputation scores were obtained from the AR2 field of the SNP-STR reference haplotype panel. **(b) Distribution of absolute spliceSTR effect sizes stratified by imputation quality.** Each box plot shows the distribution of absolute spliceSTR effect sizes (*y*-axis) across bins of imputation score (AR2 field) (*x*-axis) for the dataset (imputed STRs from HBCC dataset and directly genotyped STRs from four GTEx tissues) indicated in the panel title. Horizontal lines show median values, boxes span from the 25th percentile (Q1) to the 75th percentile (Q3). Whiskers extend to $Q1 - 1.5 \times IQR$ (bottom) and $Q3 + 1.5 \times IQR$ (top), where IQR is the interquartile range (Q3–Q1) and points indicate outlier STRs. The number of STRs considered (all spliceSTRs with FDR<5% and for which the spliceSTR is included in the SNP+STR reference panel) for each dataset is labeled under the panel title. For STRs that are significantly associated with multiple splice events, STR effect sizes were uniquified by choosing the spliceSTR with the best *P*-value.

Supplementary Figure 11

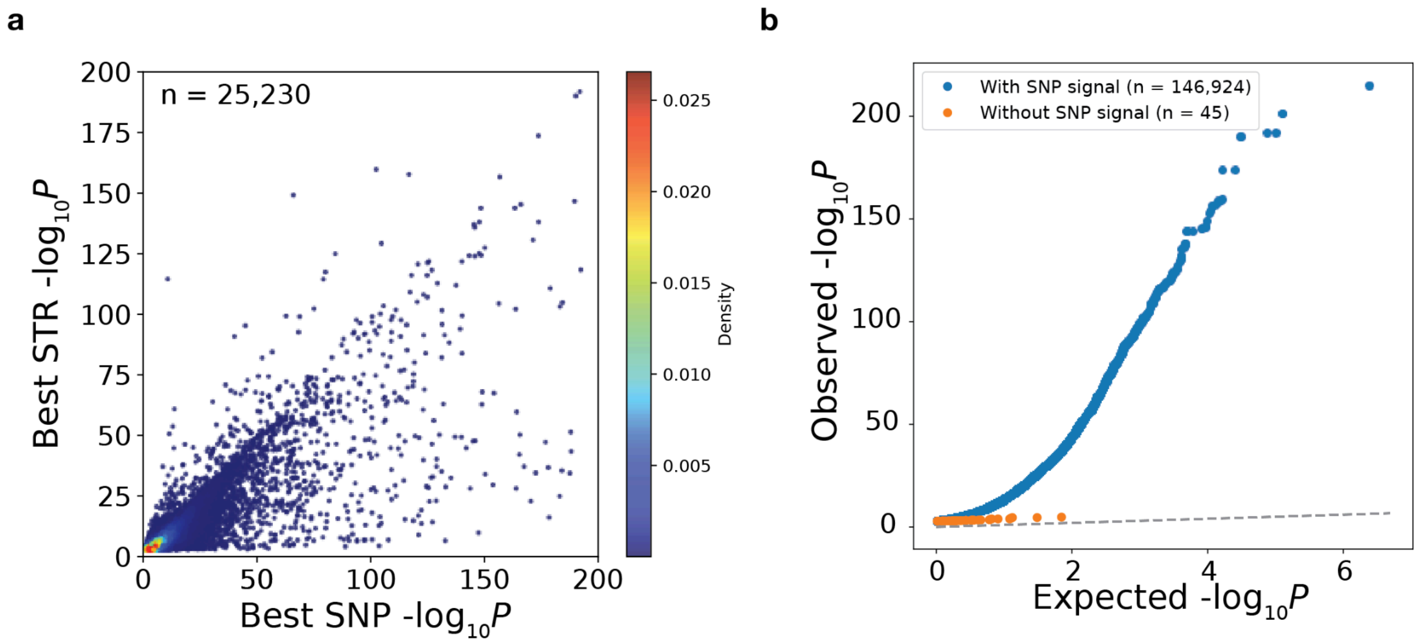


**Comparison of P-values from SNP and STR splicing association tests. (a) Comparison of the best SNP vs. best STR P-value for each splice event.** Color denotes the density of points in each region. **(b) Comparison of *P*-values between spliceSTRs with or without SNP signals detected.** The quantile-quantile plot compares the observed $-\log_{10}$ *P*-values (*y*-axis) vs. the expected $-\log_{10}$ *P*-values (*x*-axis) for spliceSTRs. SpliceSTRs were grouped by whether their associated splice events also exhibit significant SNP signals (blue=events that also have a SNP signal, orange=events that only have an STR signal). A total of 45 spliceSTRs were associated with 44 unique splicing events for which no SNP signal was detected.

### Supplementary Figure 12

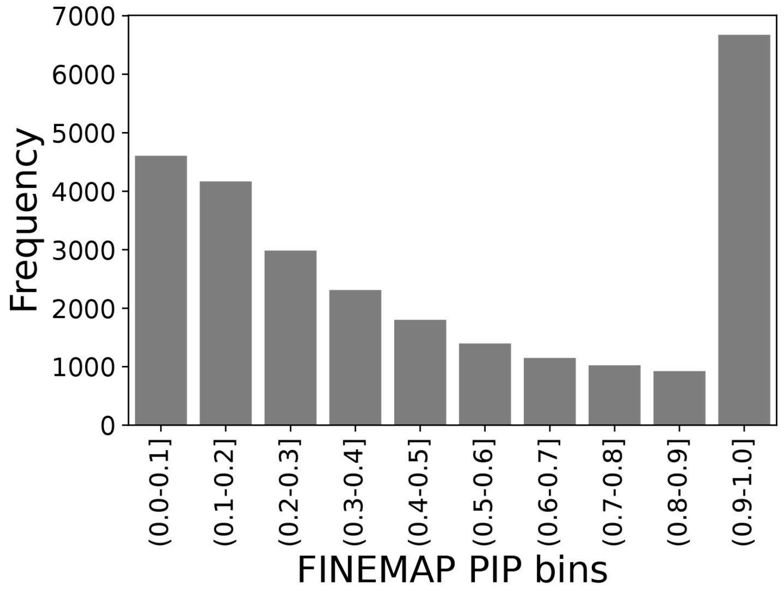


**Distribution of the top FINEMAP PIPs across all splice events.** The *x*-axis represents the bins of top PIPs for any variant (SNP or STR) for each splice event across all events tested. For 64.60% (n=15,866) of splicing events analyzed using FINEMAP, no single variant (SNP or STR) reached a threshold of PIP>0.5.

### Supplementary Figure 13

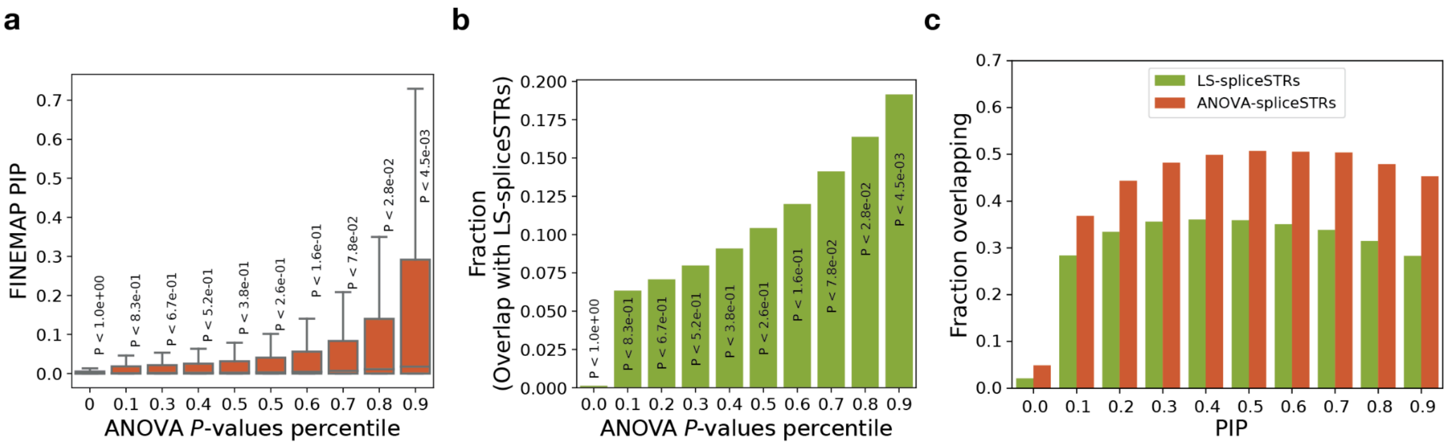


**Concordance of fine-mapping results across methods. (a) Distribution of FINEMAP PIPs for spliceSTRs thresholded by ANOVA *P*-values.** The *x*-axis represents deciles of ANOVA *P*-values (from weaker to stronger). The *y*-axis represents FINEMAP PIPs for spliceSTRs with ANOVA *P*-values below the threshold used to define each bin. Thresholds for each bin are labeled on top of each box. **(b) Fraction of spliceSTRs that are LS-spliceSTRs thresholded by ANOVA *P*-values.** The *y*-axis gives the fraction of spliceSTRs with ANOVA *P*-values below the threshold used to define each bin that are also LS-spliceSTRs. **(c) Overlap between FM-spliceSTRs and candidate spliceSTRs identified by Lasso or ANOVA.** The *x*-axis represents FINEMAP PIP thresholds used to group spliceSTRs. The *y*-axis represents the fraction of spliceSTRs with PIP above the thresholds and passing each alternative method (green=Lasso, red=ANOVA).

### Supplementary Figure 14

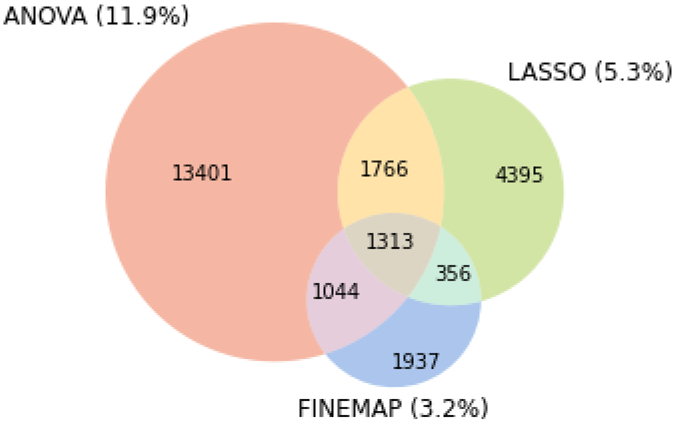


**Overlap of fine-mapped spliceSTRs across methods.** Venn diagram showing the overlap between three fine-mapping approaches: ANOVA (salmon), Lasso (green), and FINEMAP (blue). The percentages in parentheses indicates the fraction of spliceSTRs (out of all spliceSTRs passing FDR 5%) passing each method (ANOVA: 11.9%, Lasso: 5.3%, FINEMAP: 3.2%).

### Supplementary Figure 15

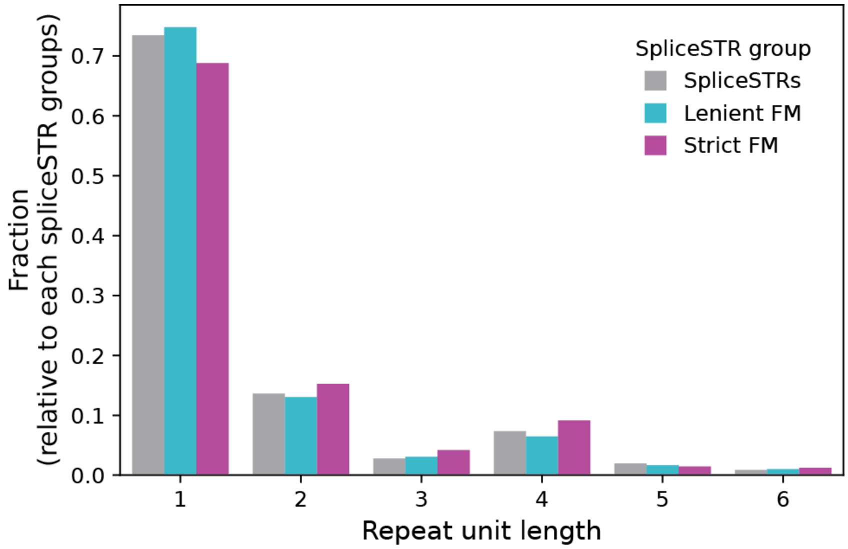


**Distribution of repeat unit length of STRs across spliceSTR sets.** The x-axis gives the repeat unit length (bp). The y-axis gives the fraction of all (gray), leniently fine-mapped (cyan), and strictly fine-mapped (purple) spliceSTRs for which the STR has each repeat unit length (bp).

### Supplementary Figure 16

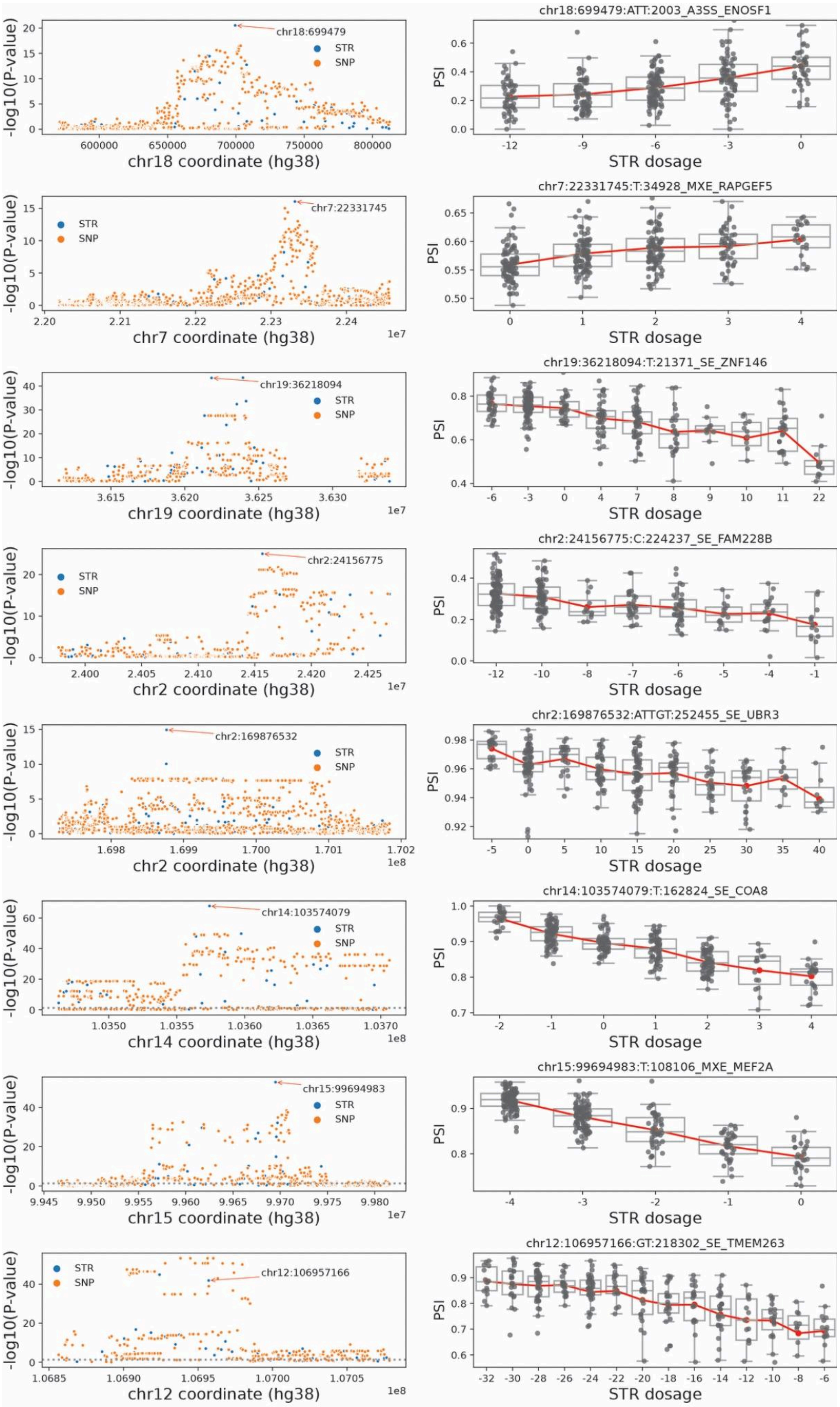

**Additional examples of fine-mapped spliceSTRs.** Each row shows a different spliceSTR for which at least two fine-mapping methods identified a candidate causal STR. Left panels show associations of all tested variants with PSI of the target splice event (blue=STRs, orange=SNPs). The candidate STR is indicated with a red arrow. Genomic coordinates are based on the hg38 reference assembly. Right panels show STR dosage (*x*-axis) vs. PSI (*y*-axis) for the target event. Note the *x*-axis ticks are evenly spaced and are in some cases not proportionally spaced according to STR dosage values.

### Supplementary Figure 17

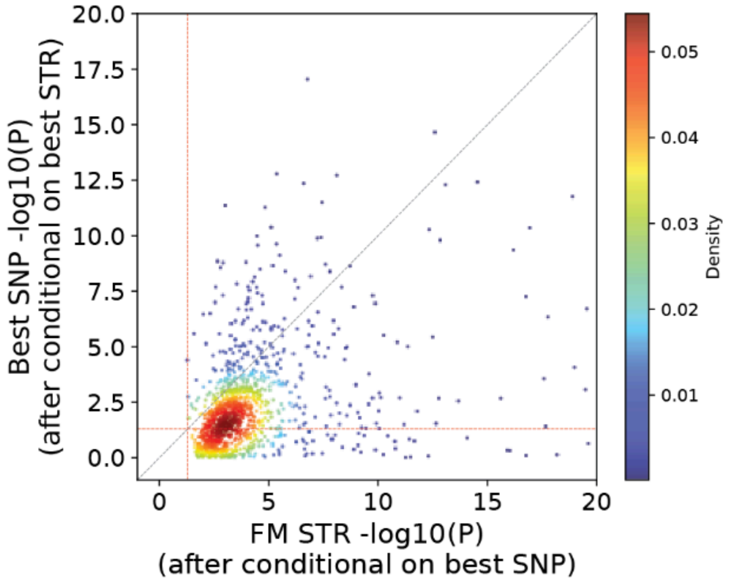


**Comparison of the best SNP vs. strictly fine-mapped STR P-value for each splice event from conditional analysis.** The *x*-axis shows the -log10 P-value of the fine-mapped STR after conditioning on the best SNP of the corresponding splice event. The *y*-axis shows the -log10 P-value of the best SNP after conditioning on the strictly fine-mapped STR of the corresponding splice event. The horizontal and vertical red dash-lines denote the nominal P-value threshold of 0.05. Color denotes the density of points in each region. The grey dash-line gives the x=y diagonal.

### Supplementary Figure 18

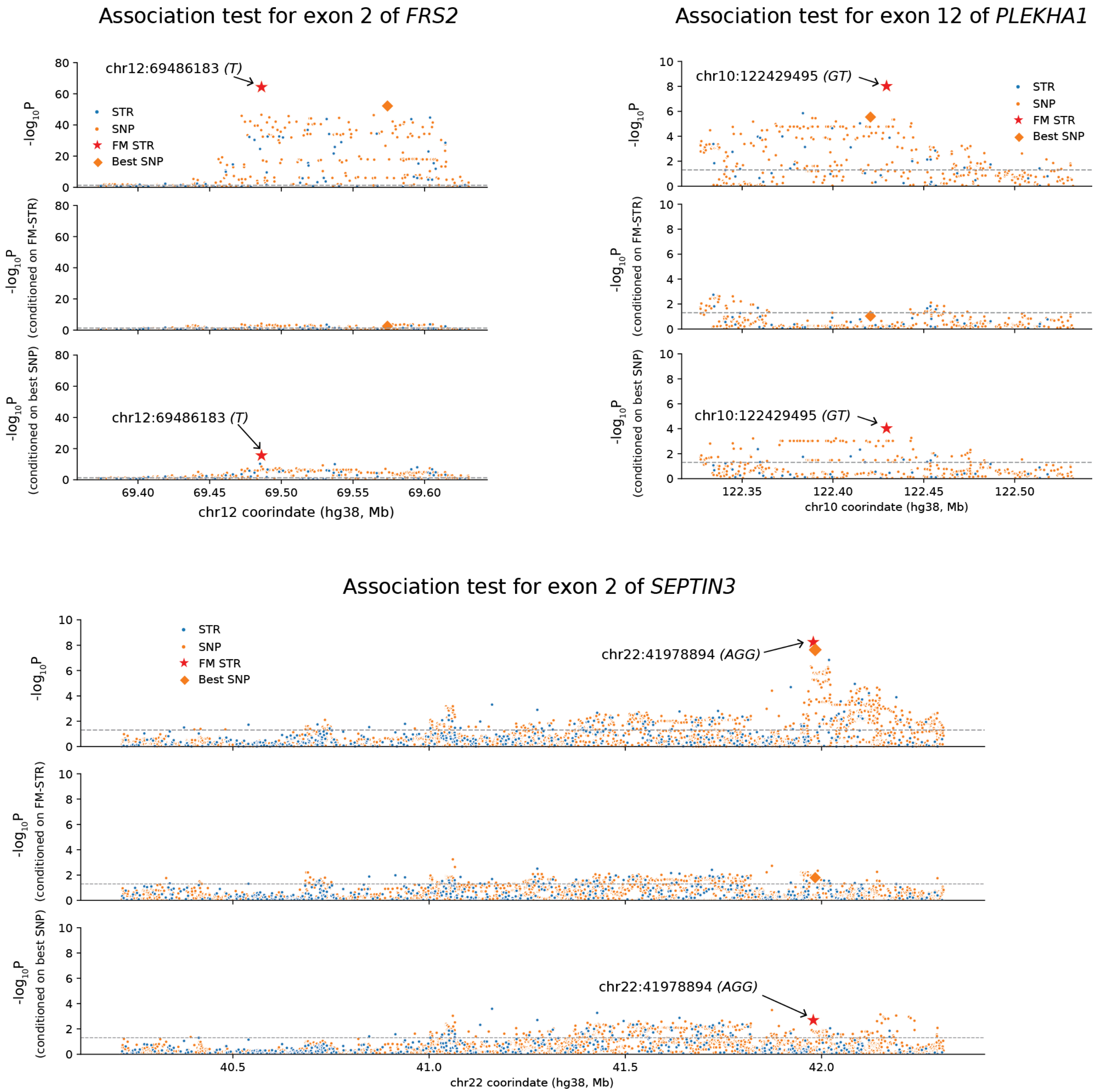


**Examples of splice event association signals with bidirectional conditional analysis.** Each plot shows the association signals for variants associated with the splice event indicated in the title. The top panel shows the unconditioned association results, the middle panel shows the association results after conditioning on the strictly fine-mapped STR, and the bottom panel shows the association results after conditioning on the lead SNP. The *x*-axis of each plot represents the genomic coordinates on the hg38 reference. Blue dots represent STRs, orange dots represent SNPs, red star dots represent strictly fine-mapped STRs and orange diamonds represent best SNPs. The *y*-axis of each dot represents -log10 *P*-values from the association test. The arrow points to the fine-mapped spliceSTR. The horizontal grey dash-line denotes the nominal *P*-value threshold at 0.05.

### Supplementary Figure 19

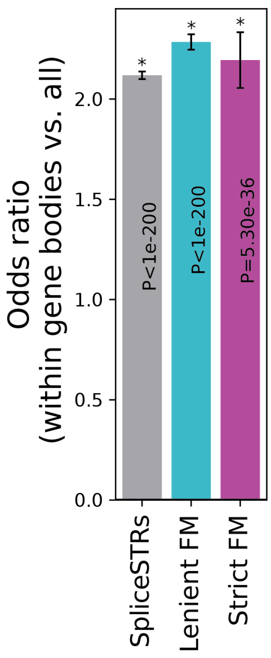


**Fine-mapped spliceSTRs are enriched within gene transcription regions.** The *y*-axis denotes the odds ratios comparing spliceSTRs or fine-mapped spliceSTRs within gene bodies to all tested STRs in each category. The *x*-axis represents the category of spliceSTR (grey=all spliceSTRs, cyan=lenient fine-mapped spliceSTRs, purple=strictly fine-mapped spliceSTRs). Error bars represent ± 1 s.e. Asterisks indicate statistically significant enrichment or depletion (two-tailed Fisher's exact test, Bonferroni corrected $P<0.05$) for each spliceSTR category.

### Supplementary Figure 20

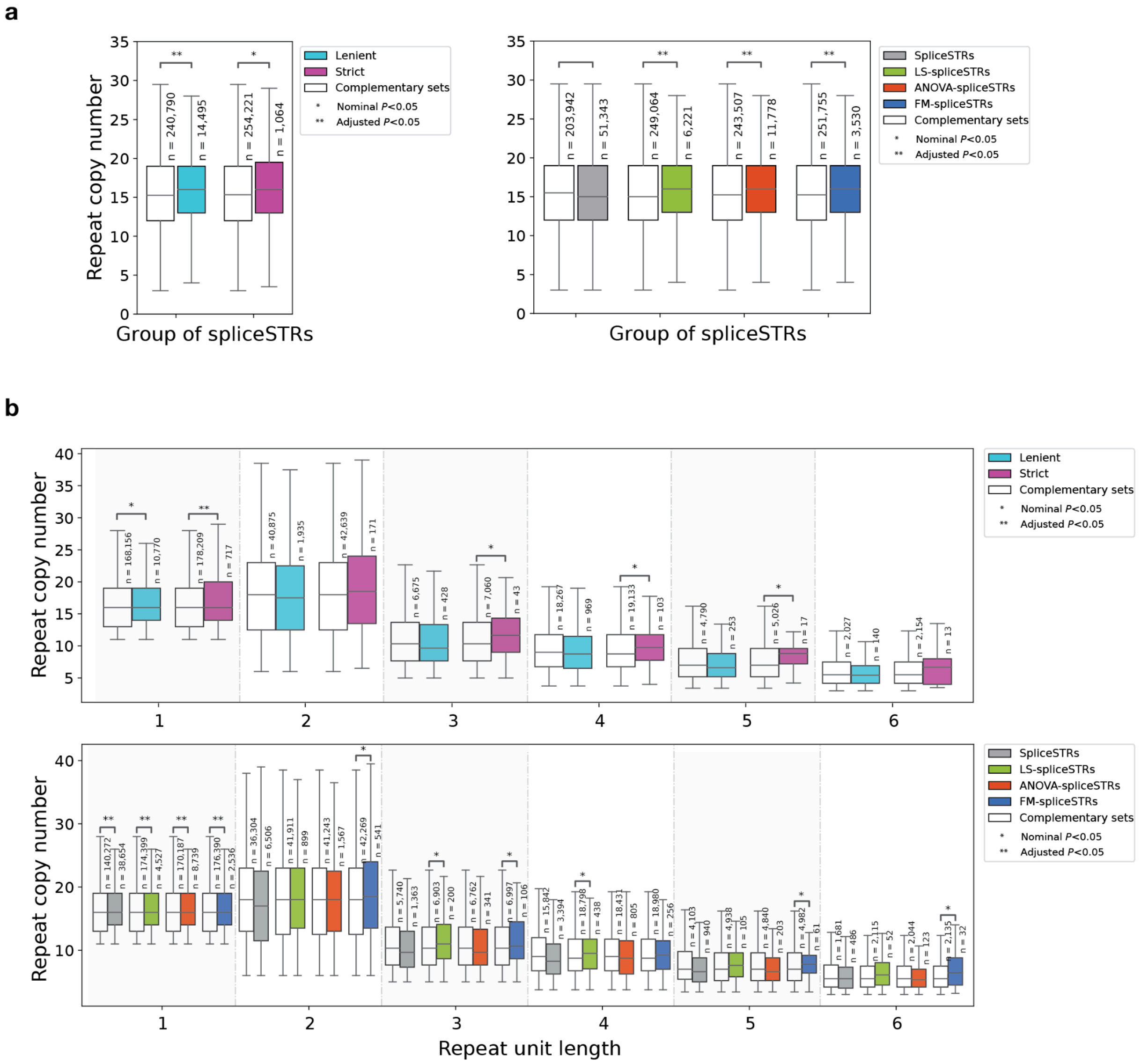


**Comparison of repeat copy numbers between fine-mapped and non–fine-mapped spliceSTRs across different repeat unit lengths and fine-mapping methods. (a) Comparison of repeat copy numbers between fine-mapped and non–fine-mapped spliceSTRs across different fine-mapping methods.** The *y*-axis shows the distribution of repeat copy numbers (based on hg38) within each group. **(b) Comparison of repeat copy numbers stratified by repeat unit length. The *x*-axis represents the repeat unit length.** The *y*-axis shows the distribution of repeat copy numbers (based on hg38) within each group. Colors indicate different fine-mapping methods, with "Complementary sets" in white denoting all tested spliceSTRs that were not fine-mapped using the method being compared to. Alternating shaded backgrounds separate repeat unit length categories. Asterisks denote statistical significance (two-sided Mann–Whitney U test) between each pair of distributions.

### Supplementary Figure 21

**a**

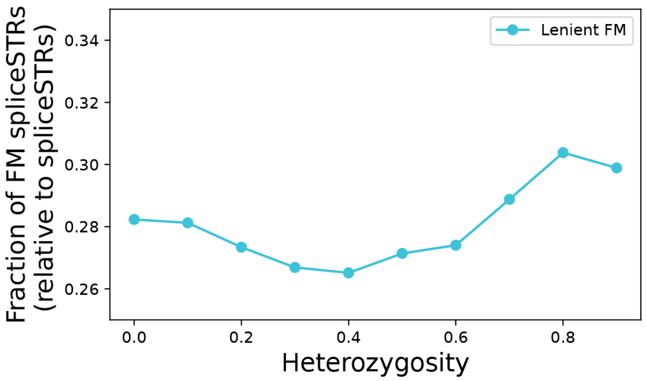


**b**

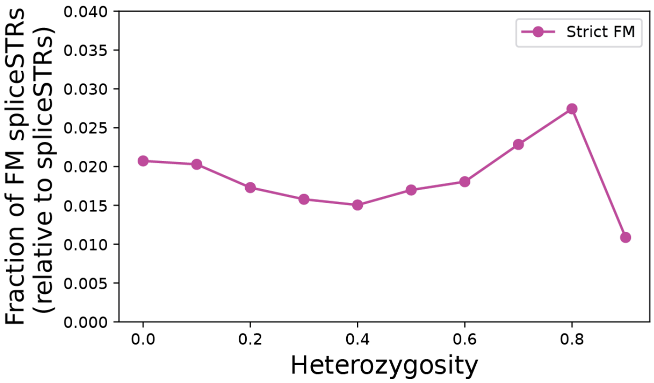


**Fraction of fine-mapped spliceSTRs across heterozygosity thresholds.** The *x*-axis denotes the minimum heterozygosity threshold to include an STR in our analysis. The *y*-axis shows the fraction of all STRs above that heterozygosity threshold that were significantly associated with at least one splice event that are also fine-mapped spliceSTRs (**a**: leniently fine-mapped spliceSTRs; **b**: strictly fine-mapped spliceSTRs).

### Supplementary Figure 22

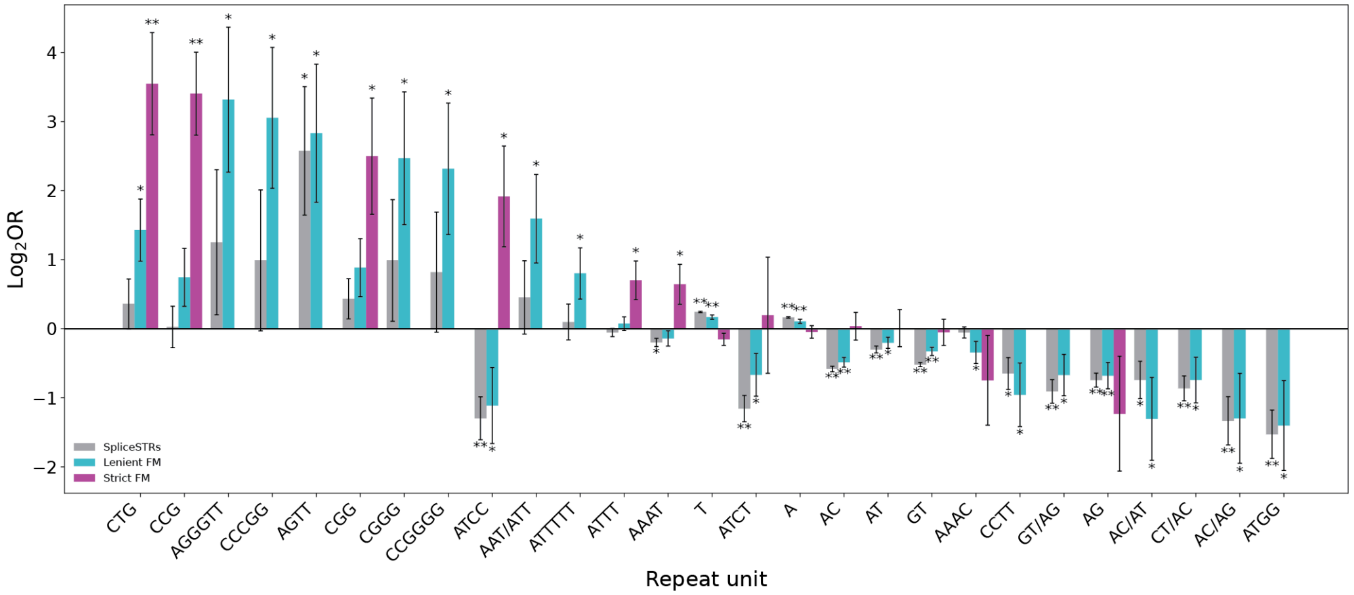


**Comparison of repeat unit enrichment across different spliceSTR categories.** The *x*-axis shows all repeat units for which there are at least 3 unique spliceSTRs for the corresponding spliceSTR group and the unit was significantly enriched. The *y*-axis denotes the $\log_2$ odds ratios comparing spliceSTRs or fine-mapped spliceSTRs to all tested STRs in each category. Error bars represent ± 1 s.e. Asterisks denote repeat units that are significantly enriched or depleted in each spliceSTR category (based on two-tailed Fisher exact *P*-value). Bars are colored by the spliceSTR category (grey=spliceSTRs, cyan=leniently fine-mapped spliceSTRs, purple=strictly fine-mapped spliceSTRs).

### Supplementary Figure 23

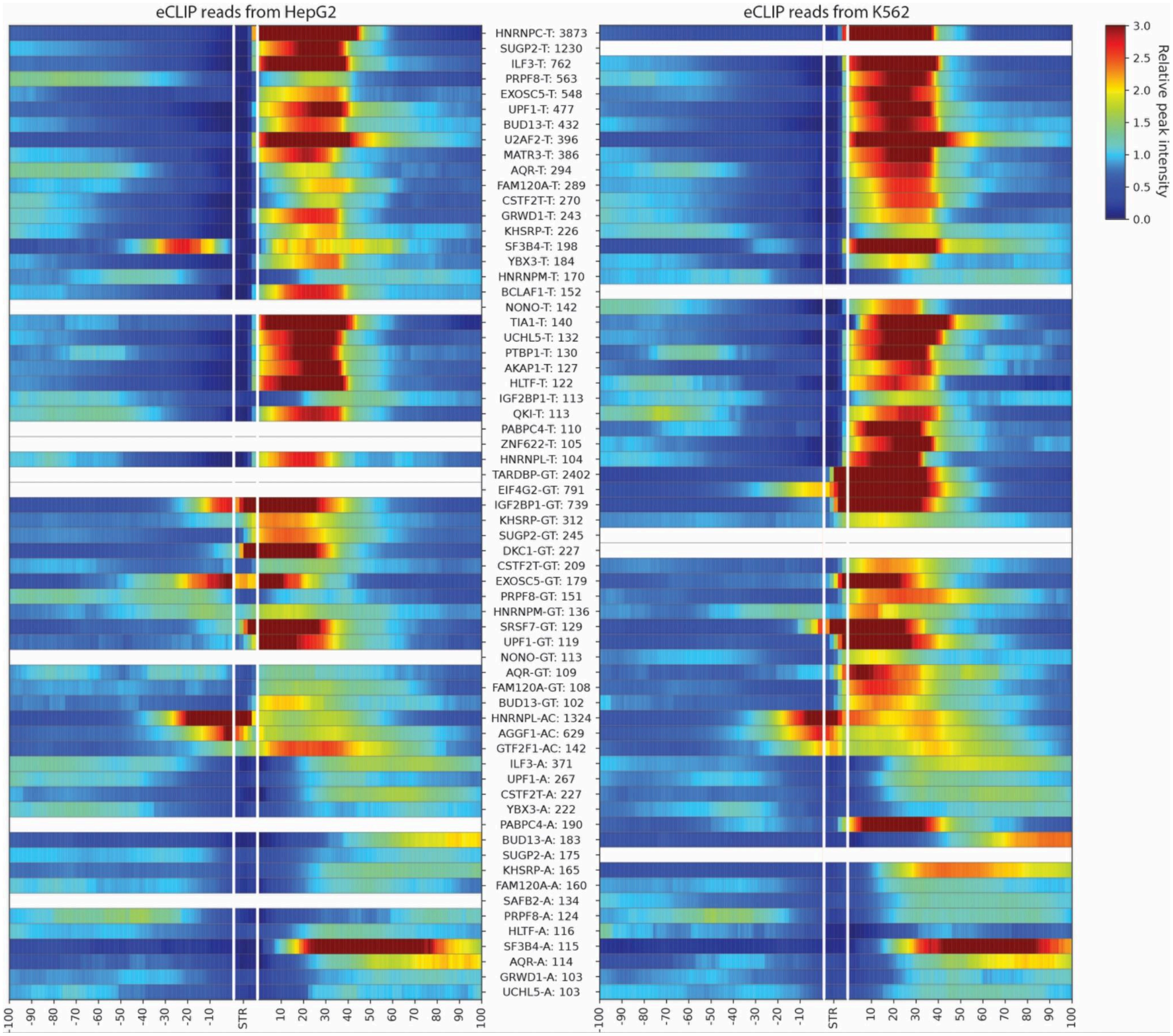


**Heatmap plot showing aggregated RBP binding profiles overlapping tested STRs stratified by repeat unit in HepG2 and K562 cell lines.** The *x*-axis represents the strand-specific distance from RNA-binding protein (RBP) binding peaks to their overlapping STR, with the STR centered at position zero. Negative values indicate upstream positions, while positive values indicate downstream. The *y*-axis lists RBP–repeat unit pairs along with the total number of overlapping repeat loci used in the analysis. Each row displays the aggregated RBP binding signal derived from the corresponding bigWig file. For each RBP–repeat unit pair, binding signals at individual loci were first normalized by the average signal of that locus, and the mean normalized signal across all loci was then used for plotting.

### Supplementary Figure 24

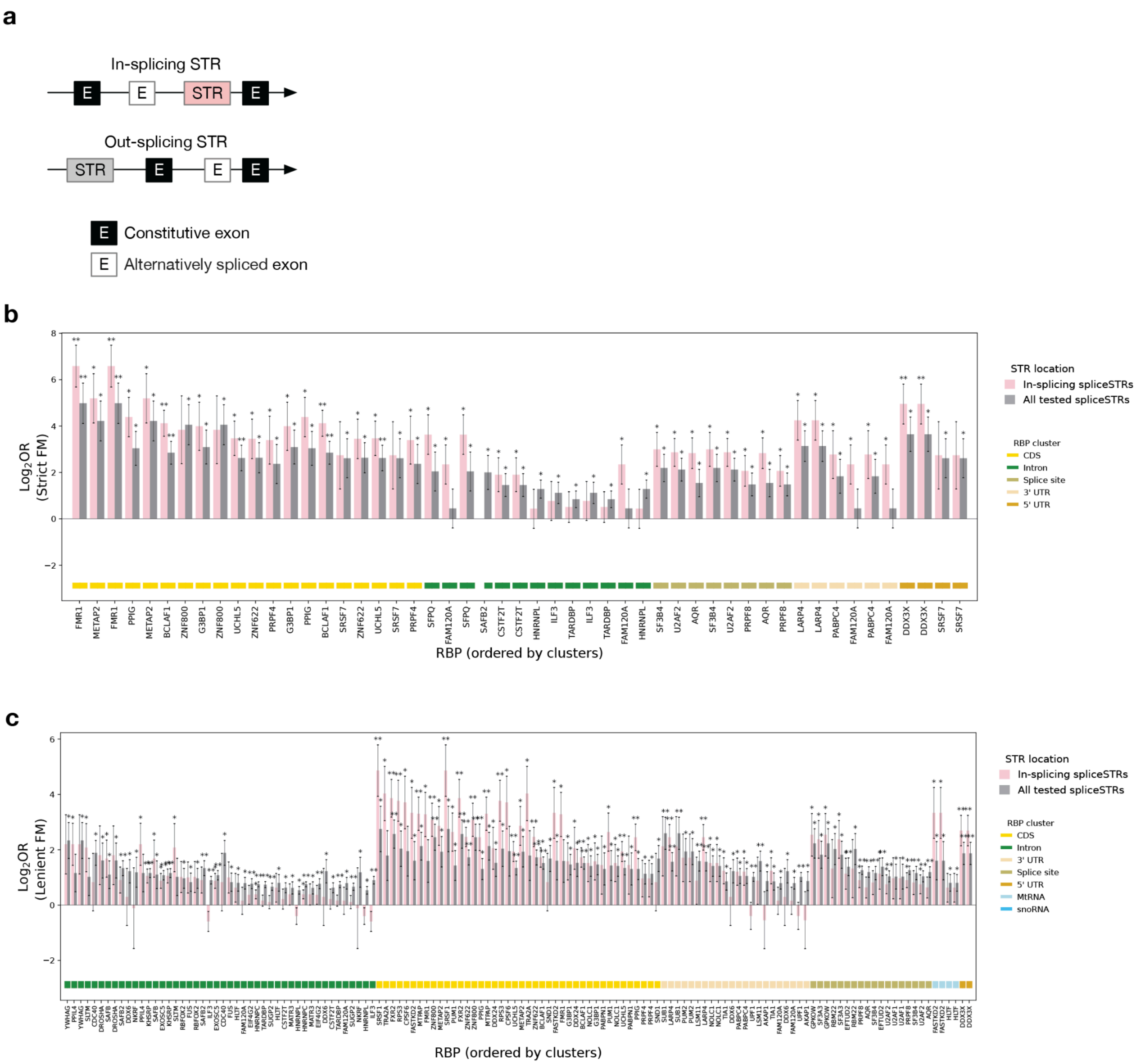


**Comparison of enrichment for overlapping of RBP binding sites stratified by the location of spliceSTR in two fine-mapped spliceSTR sets. (a) Schematic defining spliceSTRs by their position relative to associated splice events. (b) Comparison of RBP binding sites enrichment between in-splicing and out-splicing STRs for leniently and (c) strictly fine-mapped spliceSTRs.** Compared to all tested STRs, RBP enrichments are overall higher when we restrict analysis to STRs that are located within the constitutive exons of the splicing events. Compared to the leniently fine-mapped set, this trend is stronger for the strictly fine-mapped spliceSTRs. The *x*-axis shows all tested RBPs whose binding sites overlap at least three unique spliceSTRs from the corresponding fine-mapping group, as defined by Boyle et al.[4]. The y-axis represents the $\log_2$ odds ratios comparing fine-mapped spliceSTRs with all tested STRs within each STR location category for each RBP. Error bars indicate ±1 s.e. Asterisks denote RBPs for which binding sites are significantly enriched or depleted in each spliceSTR category (two-tailed Fisher's exact test, * indicates nominal $P<0.05$, ** indicates adjusted $P<0.05$). Bars are colored by spliceSTR location category (grey=all tested spliceSTRs; pink=in-splicing spliceSTRs).

### Supplementary Figure 25

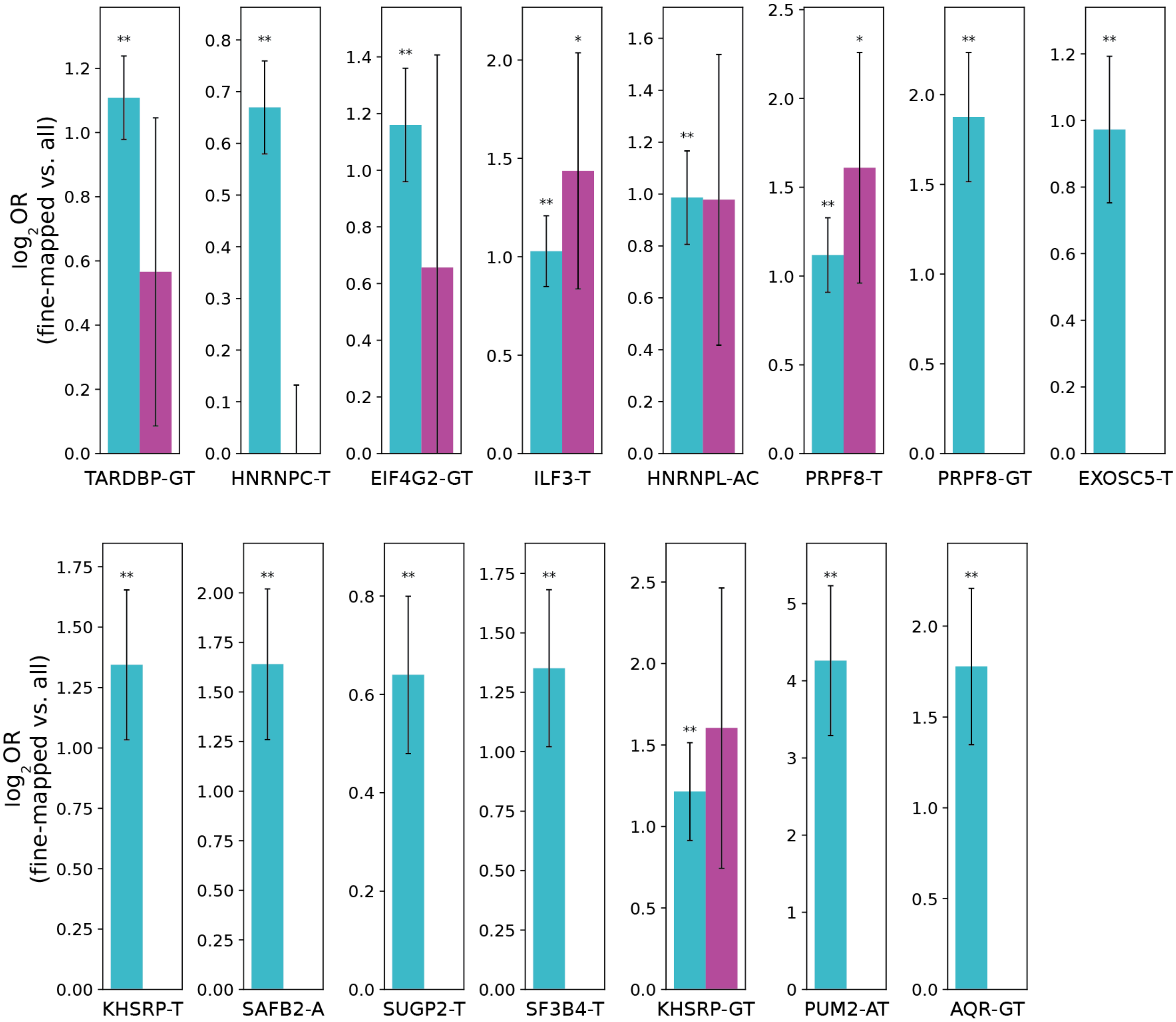


**Additional examples of enriched repeat unit-RBP pairs among fine-mapped spliceSTRs.** For each panel, the *x*-axis label represents the RBP-repeat unit pairs for which binding sites obtained from Boyle et al.[4] overlap at least 3 unique leniently fine-mapped spliceSTRs. The *y*-axis denotes the $\log_2$ odds ratio comparing each fine-mapped spliceSTR set (cyan=leniently fine-mapped spliceSTRs, purple=strictly fine-mapped spliceSTRs) to all tested STRs in each repeat unit-RBP pair. Error bars represent ± 1 s.e. Asterisks indicate statistically significant enrichment or depletion (two-tailed Fisher's exact test, * indicates nominal $P$<0.05, ** indicates adjusted $P$<0.05) for each group.

Supplementary Figure 26

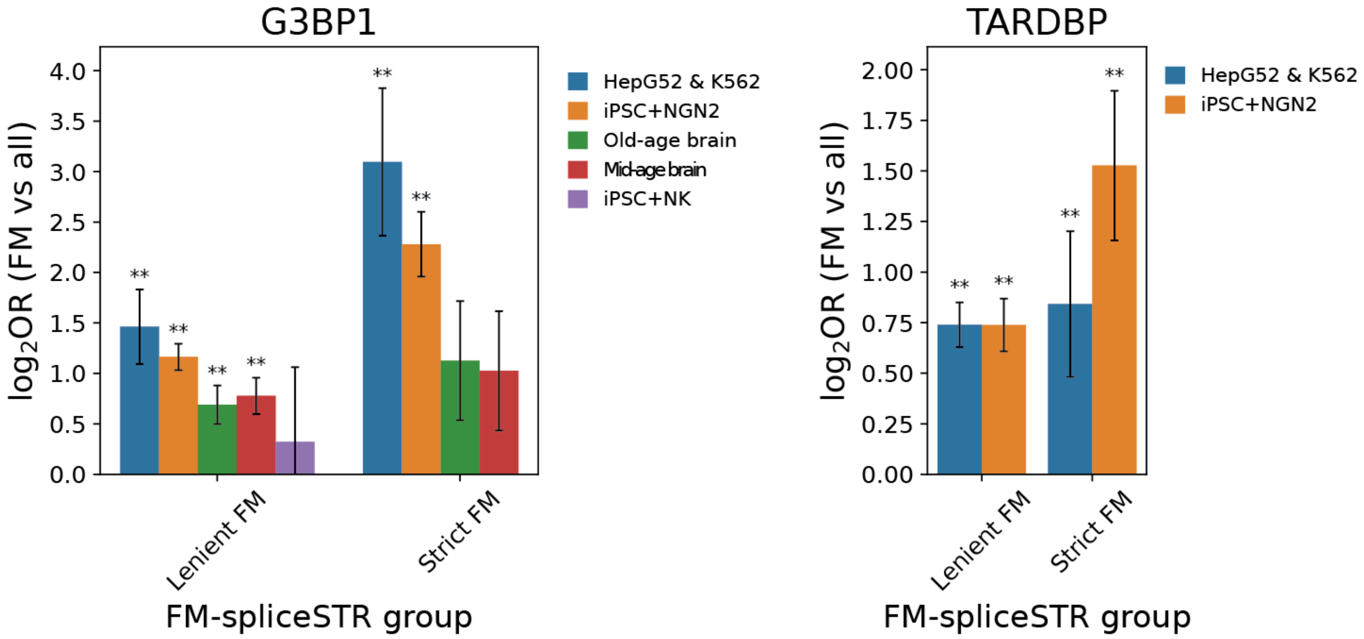


**Comparison of RBP binding site enrichment among fine-mapped spliceSTRs between non-brain and brain-related datasets for G3BP1 and TARDBP**. The *x*-axis indicates the fine-mapped spliceSTR groups, and the *y*-axis shows the log2 odds ratios for enrichment of RBP binding sites (obtained from Boyle et al. and Rhine et al.) overlapping with each fine-mapped spliceSTR group relative to all tested STRs. Results are shown for the RBP indicated in each panel title across different eCLIP datasets: HepG2 and K562 cell lines (HepG52 & K562: blue), human NGN2-induced iPSC-derived neuron cells (iPSC+NGN2: orange), old-age human frontal cortex (old-age brain: green), mid-age human frontal cortex (mid-age brain: red), and human iPSC-derived neurons (NK condition) (iPSC+NK: purple). For each RBP, only datasets in which its binding sites overlapped at least three unique fine-mapped spliceSTRs are shown. Error bars represent ± 1 s.e. Asterisks indicate statistically significant enrichment or depletion (two-tailed Fisher's exact test, * indicates nominal *P*<0.05, ** indicates adjusted *P*<0.05) for each group.

### Supplementary Figure 27

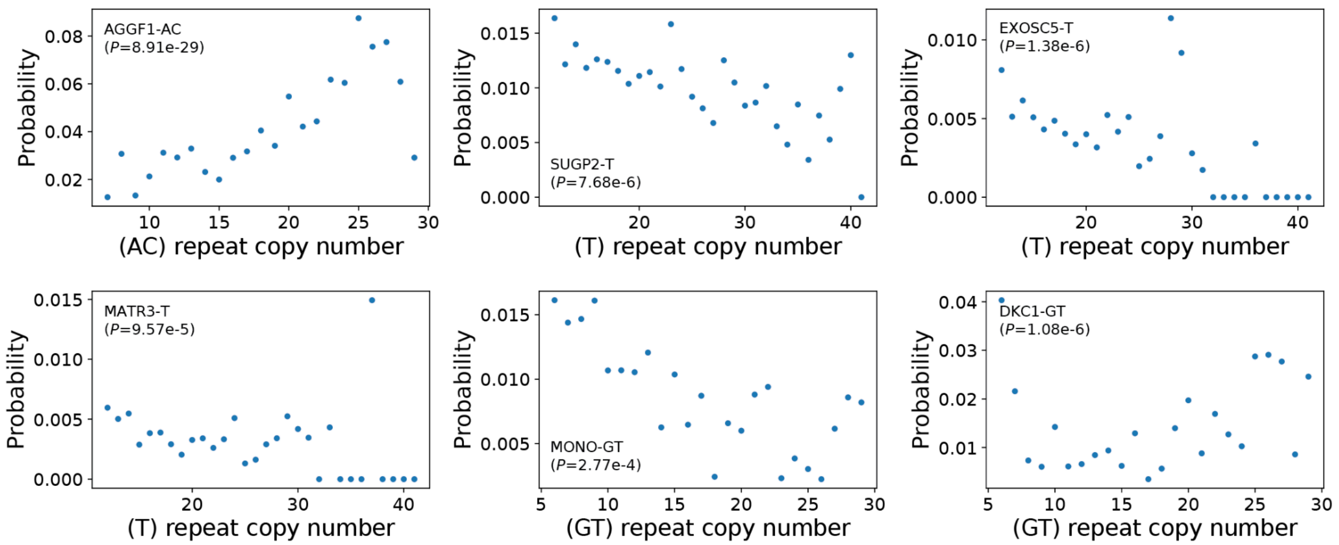


**Additional examples of significant associations between RBP binding probability and repeat unit length.** The *x*-axis represents the repeat unit copy number. The *y*-axis shows the probability of RBP binding across all STRs genome-wide of that length. *P*-values from the logistic regression are labeled in each plot.

### Supplementary Figure S28

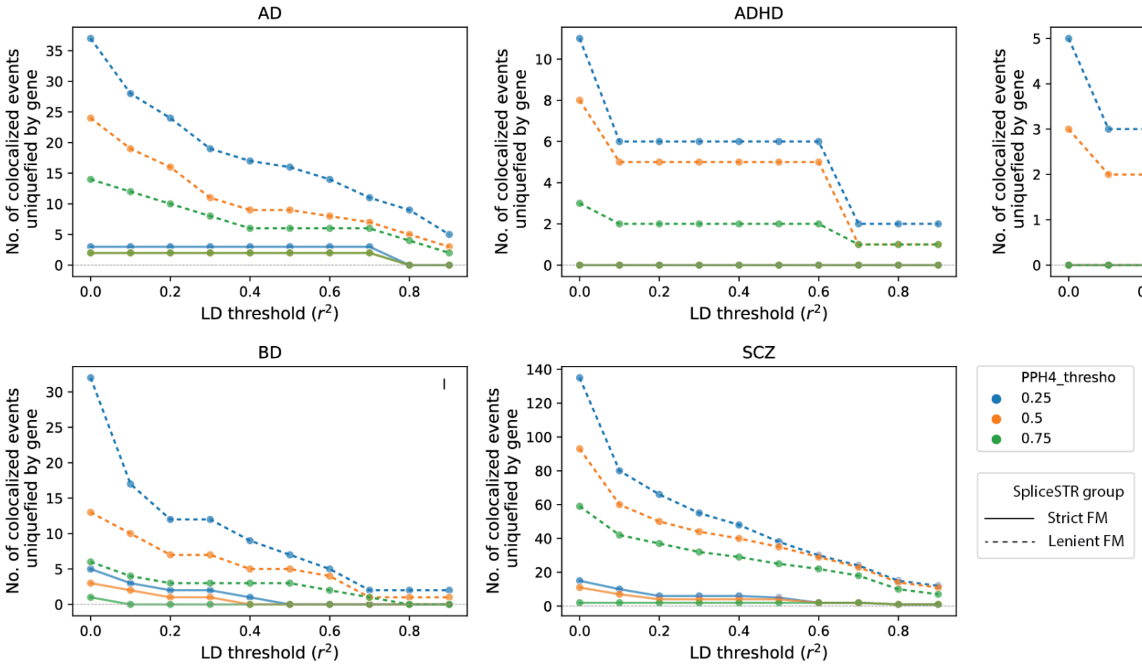


**Evaluation of the impact of PPH4 and LD $r^2$ thresholds on the number of spliceSTRs co-localized with GWAS signals identified.** Each panel is for a different trait (AD=Alzheimer's Disease, ADHD=attention deficit/hyperactivity disorder, ASD=autism spectrum disorder, BD=bipolar disease, SCZ=schizophrenia). In each panel, the *x*-axis gives the LD $r^2$ threshold and the *y*-axis gives the number of co-localized spliceSTRs identified, uniquified by gene. Color denotes the PPH4 threshold (blue=25%, orange=50%, green=75%). Solid lines denote strictly fine-mapped spliceSTRs and dashed lines denote leniently fine-mapped spliceSTRs.

### Supplementary Figure 29

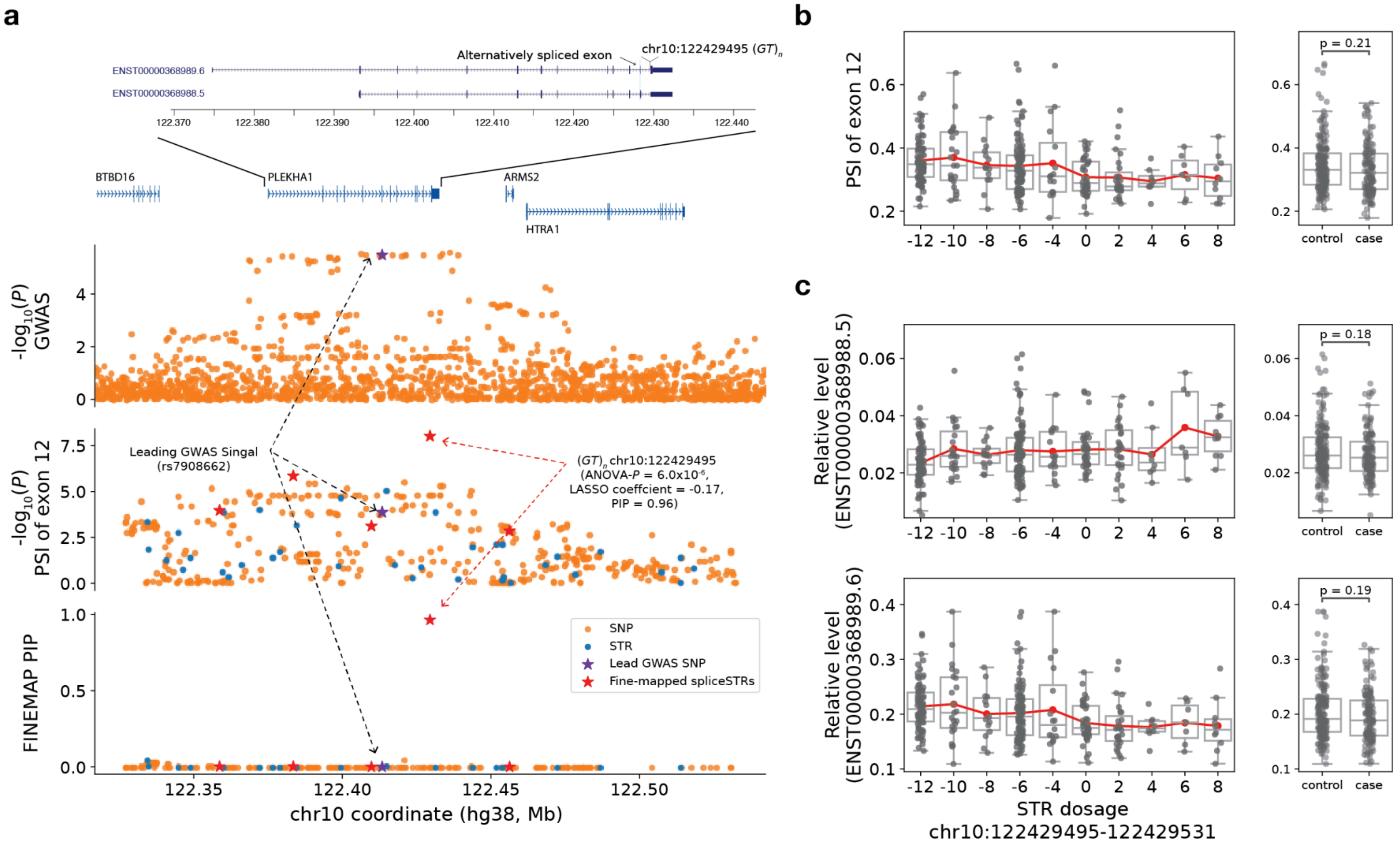


**A fine-mapped spliceSTR associated with skipping of exon 12 of *PLEKHA1* colocalizes with a GWAS signal for Alzheimer's disease. (a) A fine-mapped splice event that colocalizes with an Alzheimer's disease GWAS signal in *PLEKHA1*.** The left panels, from top to bottom, provide genomic annotations, including a zoomed-in view to show *PLEKHA1* transcripts, -log$_{10}$(*P*-values) for Alzheimer's disease GWAS, -log$_{10}$(*P*-values) for splicing association tests, and FINEMAP posterior inclusion probabilities (PIPs) for all nominally significant SNPs and STRs associated with exon 12 skipping. The *x*-axis shows genomic coordinates based on the hg38 reference. Transcripts that are significantly associated with the spliceSTRsare shown to illustrate the differentially spliced exon. (**b**) **Association between STR dosage and the PSI of exon 12 of *PLEKHA1* (ENST00000368989.6).** In the left panel: the *x*-axis represents the STR dosage (sum of repeat lengths across both chromosome copies, in bp relative to the hg38 reference) of an (*GT*)$_n$ STR (hg38:chr10:122429495-122429531) within an intron of *PLEKHA1*. The *y*-axis represents the PSI of exon 12 (ENST00000368989.6). Each dot represents a single individual. Box plots summarize the distribution of PSI values. Horizontal lines show median values, boxes span from the 25th percentile (Q1) to the 75th percentile (Q3). Whiskers extend to Q1 − 1.5 × IQR (bottom) and Q3 + 1.5 × IQR (top), where IQR is the interquartile range (Q3–Q1). The red line shows the mean expression for each x-axis value. The right panel compares the PSI of the same exon (exon 12; ENST00000368989.6) in controls (n=194) or cases (samples identified with bipolar disorder or schizophrenia, n=142). Nominal Mann-Whitney two-sided *P*-values are labeled in the plot . (**C**) **Association of STR dosage vs. *PLEKHA1* isoforms.** The left panel shows the relative abundance of *PLEKHA1* transcripts vs. STR dosage. The right panel compares the relative abundance of the transcripts in controls (n=194) and cases (samples identified with bipolar disorders or schizophrenia, n=142).

### Supplementary Figure 30

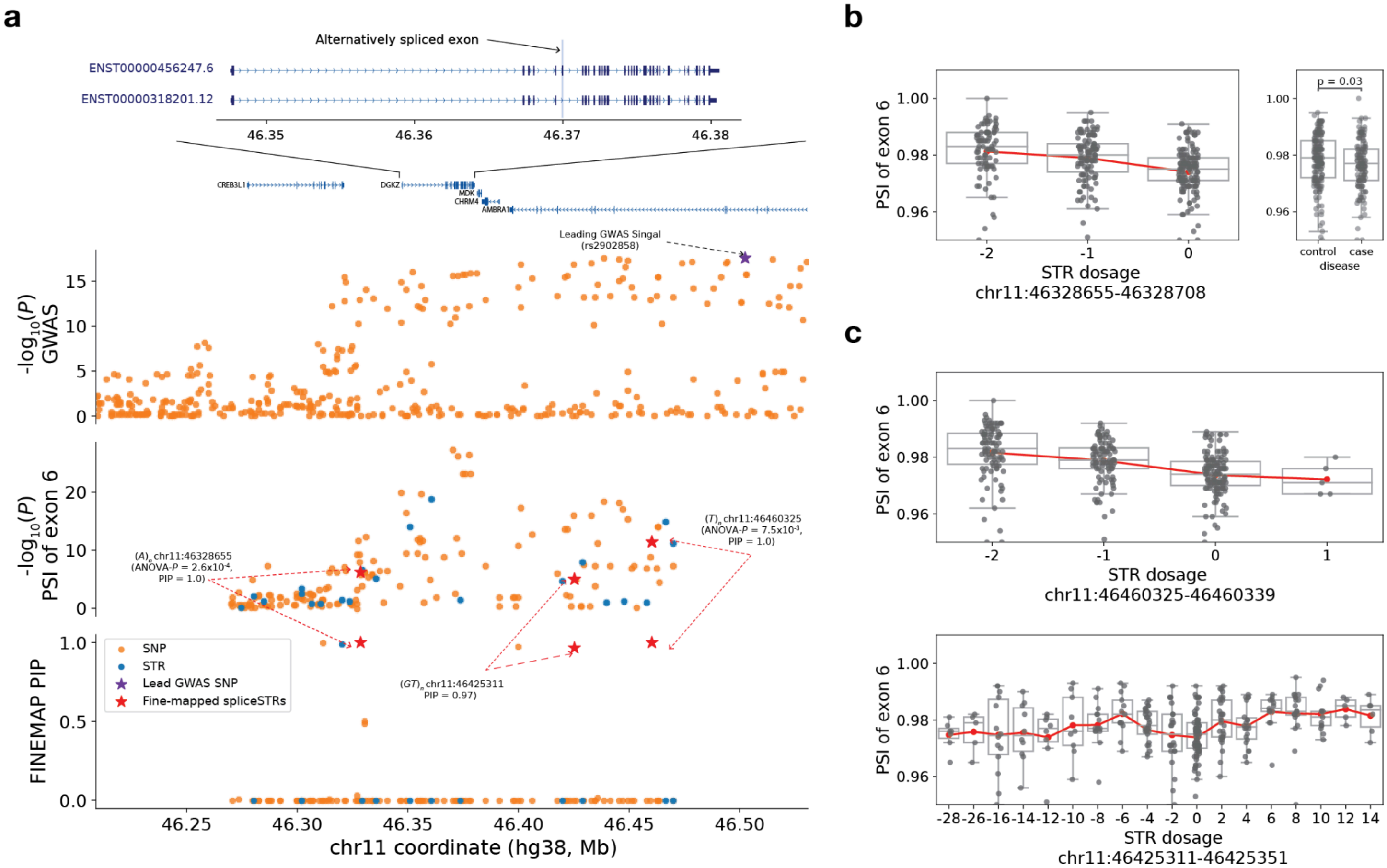


**A fine-mapped spliceSTR associated with skipping of exon 6 of *DGKZ* colocalizes with a GWAS signal for schizophrenia. (a)** A fine-mapped splice event that colocalizes with a schizophrenia GWAS signal. The left panels, from top to bottom, provide genomic annotations, including a zoomed-in view to show *DGKZ* transcripts, -$\log_{10}$(*P*-values) for schizophrenia GWAS, -$\log_{10}$(*P*-values) for splicing association tests, and FINEMAP posterior inclusion probabilities (PIPs) for all nominally significant SNPs and STRs associated with exon 6 skipping of *DGKZ (*ENST00000456247.6). The *x*-axis shows genomic coordinates based on the hg38 reference. Representative transcripts are shown to illustrate the locations of the differentially spliced exon. (**b**) Association between STR dosage (hg38:chr11:46328655-46328708) and the PSI of exon 6 of *DGKZ* (ENST00000456247.6). In the left panel: the *x*-axis represents the STR dosage (sum of repeat lengths across both chromosome copies, in bp relative to the hg38 reference) of an $(A)_n$ STR (hg38:chr11:46328655-46328708) upstream of *DGKZ*. The *y*-axis represents the PSI of exon 6 (ENST00000456247.6). Each dot represents a single individual. Box plots summarize the distribution of PSI values. Horizontal lines show median values, boxes span from the 25th percentile (Q1) to the 75th percentile (Q3). Whiskers extend to Q1 − 1.5 × IQR (bottom) and Q3 + 1.5 × IQR (top), where IQR is the interquartile range (Q3–Q1). The red line shows the mean expression for each x-axis value. The right panel compares the PSI of the same exon (exon 6; ENST00000456247.6) in controls (n=194) or cases (samples identified with bipolar disorder or schizophrenia, n=142). Nominal Mann-Whitney two-sided *P*-values are labeled in the plot . (**c**) Additional fine-mapped STRs that are associated with the PSI of exon 6 of *DGKZ* (ENST00000456247.6).

### Supplementary Figure 31

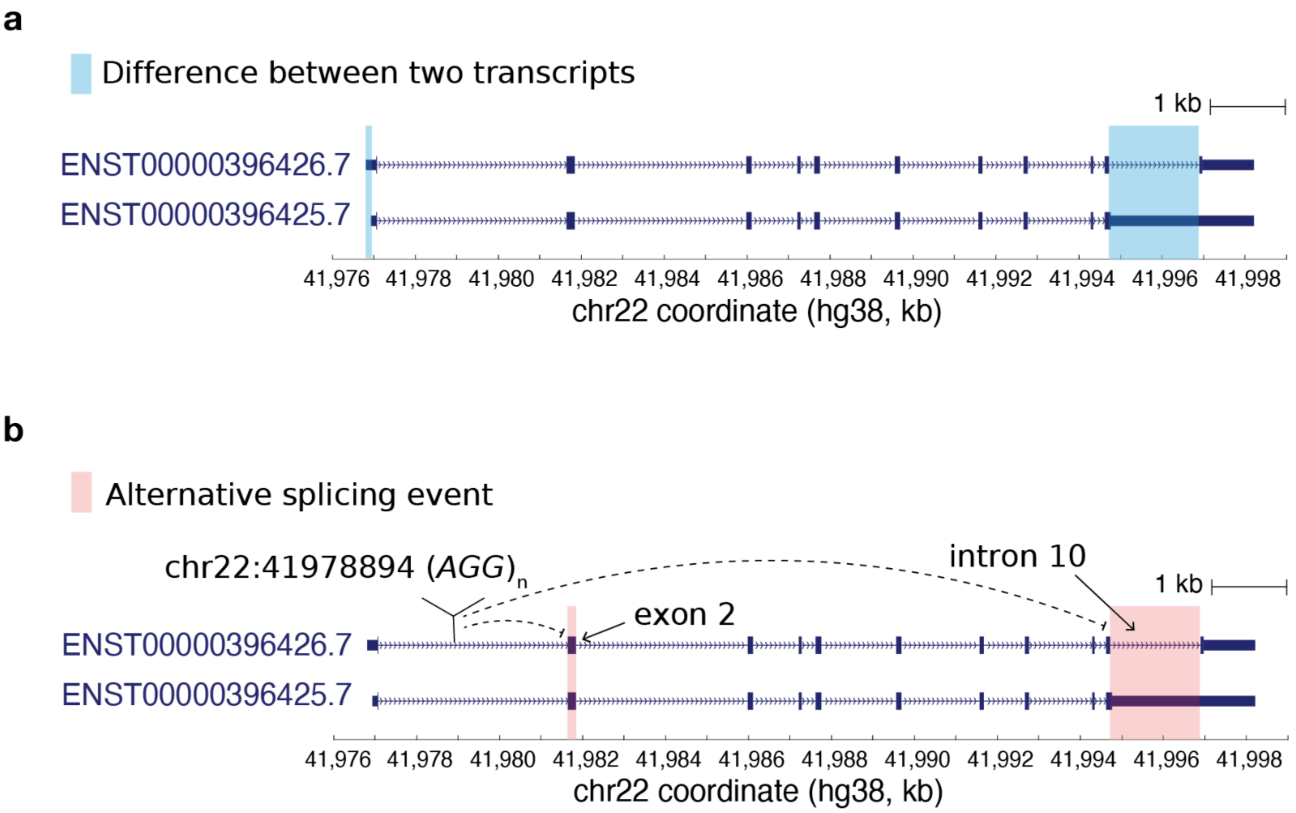


**Diagrams showing two *SEPTIN3* transcripts and the spliceSTRs that are potentially involved in their regulation. (a) Differences between two *SEPTIN3* transcripts.** Exon-intron structures of transcript ENST00000396426.7 (top) and transcript ENST00000396425.7 (bottom) are shown. The *x*-axis indicates the genomic coordinates (in kilobase) on the hg38 reference. Differences between the two transcripts are highlighted with blue boxes. **(b) SpliceSTR-associated alternative splicing events that may contribute to the formation of two SEPTIN3 transcripts.** The $(AGG)_n$ repeat (hg38:chr22:41978894-41978931) is significantly negatively associated with two splicing events, the inclusion of exon 2 and the retention of intron 10, highlighted with red boxes. The opposite effect sizes between the $(AGG)_n$ repeat (hg38 chr22:41978894–41978931) and the abundance of these two transcripts may be explained by their opposing patterns of intron 10 usage.

### Supplementary Figure 32

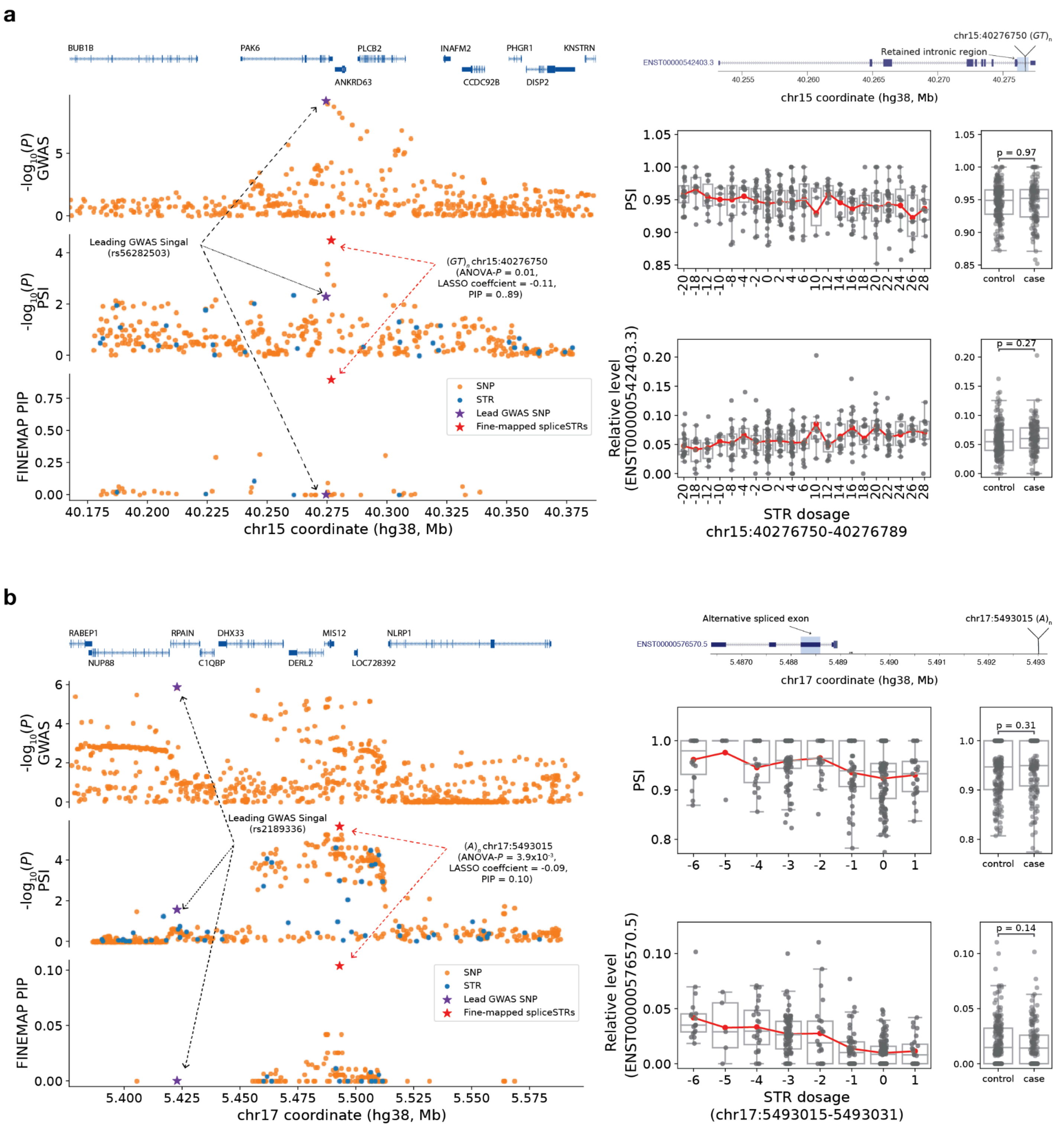


**Additional examples of fine-mapped spliceSTRs that colocalized with GWAS signals. (a) A fine-mapped spliceSTR associated with an intron retention of *PAK6* colocalizes with a GWAS signal for schizophrenia.** The left panels, from top to bottom, provide genomic annotations, -$\log_{10}$(*P*-values) for schizophrenia GWAS, -$\log_{10}$(*P*-values) for splicing association tests, and FINEMAP posterior inclusion probabilities (PIPs) for all nominally significant SNPs and STRs associated with an intron retention event of *PAK6 (*ENST00000542403.3). The *x*-axis shows genomic coordinates based on the hg38 reference. The right panels show a zoomed-in view to show *PAK6* transcripts (top panel), including the location of a retained intronic region and its fine-mapped STR, the association of STR dosage vs. the PSI of the intron retention (ENST00000542403.3, middle panel), and a *PAK6* isoform that is associated with that colocalized fine-mapped spliceSTR (bottom panel), along with box plots comparing the PSI of intron retention of *PAK6,* or the relative abundance of the transcript ENST00000542403.3 in controls (n=194) and cases (samples identified with bipolar disorders or schizophrenia, n=142). Nominal Mann-Whitney two-sided *P*-values are labeled in the plot. Each dot represents a single individual. Box plots summarize the distribution of PSI values. Horizontal lines show median values, boxes span from the 25th percentile (Q1) to the 75th percentile (Q3). Whiskers extend to Q1 − 1.5 × IQR (bottom) and Q3 + 1.5 × IQR (top), where IQR is the interquartile range (Q3–Q1). The red line shows the mean expression for each *x*-axis value. **(b) A fine-mapped spliceSTR associated with skipping of exon 3 of *MIS12* colocalizes with a GWAS signal for schizophrenia.** The left panels, from top to bottom, provide genomic annotations, -log10(*P*-values) for schizophrenia GWAS, -log10(*P*-values) for splicing association tests, and FINEMAP posterior inclusion probabilities (PIPs) for all nominally significant SNPs and STRs associated with the skipping of exon 3 of *MIS12 (*ENST00000576570.5). The *x*-axis shows genomic coordinates based on the hg38 reference. The right panels show a zoomed-in view to show *MIS12* transcripts (top panel), including the location of exon 3 and its fine-mapped STR, the association of STR dosage vs. the PSI of the skipping of exon 3 of *MIS12* (ENST00000576570.5, middle panel), and a *MIS12* isoform that associated with that colocalized fine-mapped spliceSTR (bottom panel), along with box plots comparing the PSI of exon 3 of *MIS12,* or the relative abundance of the transcript ENST00000576570.5 in controls (n=194) and cases (samples identified with bipolar disorders or schizophrenia, n=142). For both panels, transcripts which are significantly associated with the spliceSTRs are shown to illustrate the locations of the differentially spliced regions.
